# GM-CSF and M-CSF Driven Differentiation Differentially Regulates Chikungunya Virus Infection and Antiviral Responses in Human Monocyte-Derived Macrophages

**DOI:** 10.64898/2026.03.11.710213

**Authors:** Jeury Veloz, Victoria Zyulina, Sara Thannickal, Eva Chebishev, Dabeiba Bernal-Rubio, Maria del Mar Villanueva Guzman, Chenyang Wu, Estefania Valencia, Danielle Novillo, Vrushali Dhamapurkar, Laura E. Barranco, Laurence G. Webb, Rafael Fenutria, Maria Gabriela Noval, Ana Fernandez-Sesma

## Abstract

Chikungunya virus is an arthritogenic alphavirus causing debilitating joint pain in infected individuals. The mechanisms driving CHIKV-associated arthralgia is poorly understood, however, macrophages have been implicated as potential reservoirs of persistent viral material and mediators of immunopathology. Granulocyte Macrophage-Colony Stimulating Factor (GM-CSF) and Macrophage-Colony Stimulating Factor (M-CSF) are cytokines that serve as myeloid growth factors that bias macrophages toward pro-inflammatory and anti-inflammatory phenotypes, respectively. In this study, we examined how cytokine-driven macrophage differentiation via GM-CSF and M-CSF influences susceptibility to and responses against CHIKV infection *in vitro*. Using parallel donor-matched cultures of primary macrophages, we show that GM-CSF-differentiated macrophages are highly permissive to CHIKV and Mayaro virus (MAYV) infection and support robust viral replication, whereas M-CSF-differentiated macrophages are resistant to CHIKV replication and lack detectable levels of viral protein expression. Despite these differences, we observed pro-inflammatory, M1-skewing of CHIKV-infected macrophages, regardless of differentiation state. Interestingly, we observe higher production of IFNα and IP10/CXCL10 in M-CSF differentiated macrophages, suggesting that M-CSF promotes an antiviral state that restricts CHIKV infection. Stimulation of macrophages with double-stranded RNA (polyinosinic:polycytidylic acid; poly(I:C)), but not with single-stranded RNA (resiquimod, R848), recapitulated the antiviral cytokine and chemokine response induced by CHIKV infection. These findings suggest that dsRNA sensing plays a more prominent role than ssRNA sensing in driving the macrophage antiviral response to CHIKV. Together, these findings highlight macrophage differentiation as a critical determinant of CHIKV susceptibility and antiviral immunity in humans, with implications for understanding inflammatory pathogenesis during infection.

## INTRODUCTION

Chikungunya virus (CHIKV) is an old-world alphavirus of the *Togaviridae* family spread in humans through the bite of an infected *Aedes* species of mosquito. An estimated 35 million human infections occur annually, primarily affecting regions across Africa, South Asia, South America, and the Caribbean [1, 2]. Though epidemics of CHIKV mainly occur in tropical and subtropical climates, autochthonous cases have been identified in more than 110 countries worldwide, including the United States, European countries, and China, which is currently experiencing an explosive outbreak [3, 4]. Climate change threatens to expand vector distribution, potentially spreading CHIKV to immunologically naïve regions and causing large outbreaks among susceptible populations [3].

Chikungunya virus disease (CHIKVD) is associated with high rates of morbidity, with over 90% of affected individuals developing febrile illness, with symptoms including, but not limited to, rash, fever, and debilitating joint pain [5–7]. The severity and mortality of CHIKVD varies widely across epidemics, with excess deaths reported in a number of outbreaks over the last decade in Asia, South America, and the Caribbean [8]. CHIKVD can be self-limiting, however high viremia and sustained inflammation can drive disease progression to the subacute and chronic phases marked by debilitating joint pain and prolonged myalgia for months to years after initial exposure, affecting approximately 50% of infected individuals [9]. Treatment of CHIKVD is currently limited to supportive care for symptom relief, as specific antivirals are still an area of ongoing investigation and not available. Currently, a virus-like particle vaccine (VIMKUNYA) is available in the United States and Europe for high-risk individuals, with rollout to low- and middle-income countries expected to come in the future [10, 11]. A second live-attenuated vaccine, IXCHIQ, is subject to ongoing safety review, though it has also been shown to be highly protective among immunized individuals [12, 13].

Macrophages are innate immune effector cells of either embryonic tissue-resident or monocyte-derived origin that are essential for pathogen control and the coordination of tissue repair after infection [14]. In mice, depletion of macrophages led to amelioration of CHIKV-associated foot swelling but increased viremia [15], highlighting the double-edged role of macrophages in balancing the inflammatory response to infection. CHIKV-positive resident and infiltrating macrophages have been identified in tissue compartments of mice and non-human primates, including a recent study demonstrating active replication of CHIKV in joint macrophages of chronically infected mice [16–19]. In humans, clinical evidence demonstrates the presence of CHIKV antigen- and RNA-positive macrophages in the synovium of chronic and post-mortem severe CHIKVD patients, implicating macrophages as reservoirs for viral infection [20, 21].

During CHIKV infection, monocytes facilitate viral dissemination by infiltrating to distal tissues such as the liver, brain, and joint tissue [22, 23]. Monocytes can differentiate into effector innate immune cells, including macrophages, via cytokines produced by cells in the local cellular microenvironment. Human monocyte-derived macrophages (MDMs) have been previously used as an infection model for alphaviruses, demonstrating support for CHIKV and Mayaro virus (MAYV) replication and promoting a pro-inflammatory immune response through production of pro-inflammatory cytokines and type I interferons (IFNs) [24–27]. However, considering the functional and phenotypic plasticity of macrophages, little is known about how differentiation under pro-inflammatory or anti-inflammatory conditions influences their responses to CHIKV infection. Granulocyte-macrophage colony-stimulating factor (GM-CSF) and macrophage colony-stimulating factor (M-CSF) are members of the colony-stimulating factor (CSF) superfamily that are critical for the maturation of macrophages and their function [28, 29]. GM-CSF is induced under pro-inflammatory conditions and is elevated in serum of patients with severe and chronic CHIKV infection [30, 31]. M-CSF is constitutively expressed during homeostatic conditions by a diverse range of cell types including fibroblasts, monocytes, osteoclasts, endothelial cells and differentiates macrophages into an immunosuppressive phenotype [32]. In contrast to GM-CSF, M-CSF has been found to be downregulated in the serum of patients with acute CHIKVD [33, 34]. The role of CSF cytokines in CHIKV immunopathology in humans remains poorly defined, however depletion of GM-CSF with anti-GMCSF antibodies has been shown to reduce disease severity in CHIKV infected mice [35]. Additionally, GM-CSF and M-CSF have emerged as targets for immunotherapy to modulate infectious disease associated inflammatory pathology and rheumatoid arthritis, an autoimmune disease that shares clinical features with chronic CHIKVD [36–42]. Thus, elucidating how GM-CSF and M-CSF separately influence macrophage responses to CHIKV infection could reveal new immunotherapeutic avenues to curb pathogenic macrophage and myeloid cell responses.

While a direct link between GM-CSF or M-CSF signaling and CHIKV infection in macrophages has not yet been established, evidence from mouse models suggest that skewing of macrophage function towards a pro-inflammatory phenotype is crucial for reducing viremia and limiting disease severity, underscoring the central role of macrophage function in determining CHIKV disease progression [43, 44]. Given the critical role of GM-CSF and M-CSF in determining macrophage function, we sought to determine how cytokine-driven macrophage differentiation shapes susceptibility to CHIKV infection and downstream innate immune responses in human MDMs.

## RESULTS

### GM-CSF but not M-CSF directed differentiation enables susceptibility to CHIKV and MAYV infection in human MDMs

Infection and replication of CHIKV and other alphaviruses in primary human macrophages has been previously described [20, 26, 27, 45]. However, whether differentiation of MDMs with GM-CSF or M-CSF influences the susceptibility of macrophages to CHIKV infection has not been previously explored. To answer this, we isolated CD14^+^ monocytes from healthy human peripheral blood and further differentiated the monocytes with GM-CSF (GM-Mϕ) or M-CSF (M-Mϕ) for 6 days and infected them with CHIKV for up to 48 hours. First, we assessed intracellular viral replication in macrophages infected with CHIKV 181/25 or its parental strain CHIKV AF15561 by RT-qPCR (Fig. 1A-1B). Active replication of alphaviruses is dependent on successful transcription of positive-stranded genomic and subgenomic RNAs, leading to translation and production of non-structural and structural viral proteins, respectively [46]. To determine whether macrophages support active replication of CHIKV, we measured the ratio of E1 (subgenomic) to nsP1 (genomic) viral RNA transcripts produced over 48 hours of infection via standard curve, as previously done [47]. We observed significantly higher active replication of CHIKV 181/25 at 6 and 24 hours post infection (hpi) and AF15561 at 24 hpi in GM-Mϕ than M-Mϕ, suggesting GM-CSF but not M-CSF may facilitate conditions for viral replication and persistence in macrophages.

**Figure 1:**
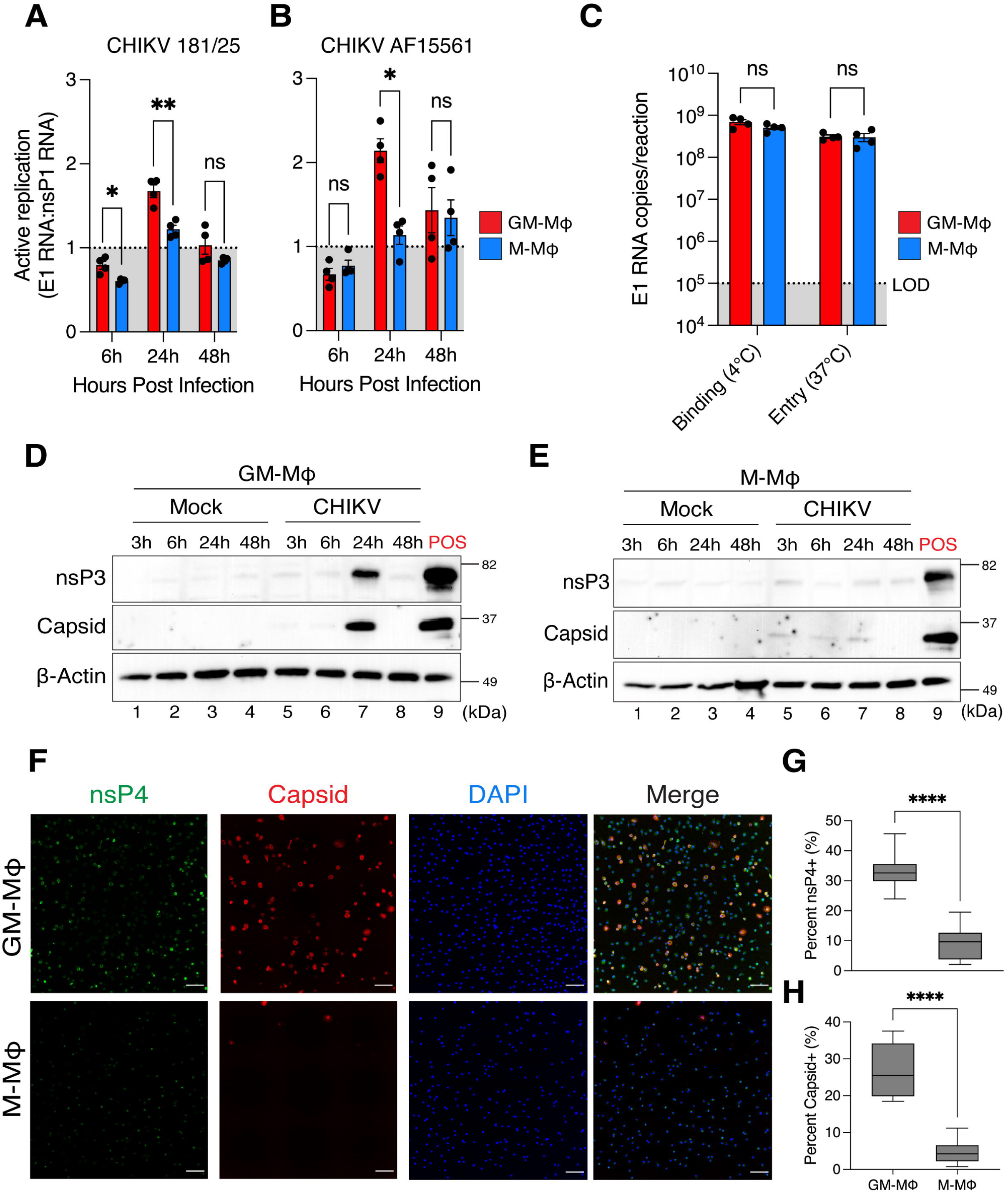
Viral infectivity and replication of Chikungunya virus in primary human macrophages. **(A-B)** Monocyte-derived macrophages were differentiated with either GM-CSF (GM-M⏀) or M-CSF (M-M⏀) for 6 days and infected with CHIKV 181/25 or CHIKV AF15661 at a multiplicity of infection (MOI) of 1.0 for 6, 24, and 48 hours. Intracellular viral replication is represented as “active replication”, which was determined by evaluating the ratio of E1 subgenomic viral RNA copies to nsP1 genomic viral RNA copies. (E1RNA:nsP1RNA =1 no replication; E1:nsP1>1 active replication) of viral RNA was determined by RT-qPCR via standard curve of known concentrations of CHIKV *in vitro* transcribed (IVT). Data represented as means ± SEM (n= 4 donors). Statistical analysis was performed via two-way ANOVA with Geisser-Greenhouse correction and Fisher’s LSD test. **(C)** Quantification of E1 viral RNA copies (log scale) in GM-M⏀ (red bars) and M-M⏀ (blue bars) collected after incubation on ice (viral binding) or incubation for 2 hours at 37°C (viral entry). Data represented as means ± SEM (n= 4 donors). Statistical analysis was performed via two-way ANOVA with Geisser-Greenhouse correction and Fisher’s LSD test. **(D-E)** Mock or CHIKV 181/25 infected (MOI=1.0) GM-M⏀ and M-M⏀ were lysed at indicated timepoints for SDS-PAGE and immunoblotting. ”POS” indicates a positive control lane of primary HFF-1 human foreskin fibroblast cells infected with CHIKV 181/25 collected at 24 hpi. **F)** GM-M⏀ and M-M⏀ were infected with CHIKV 181/25 (MOI=1.0) for 24 hours and fixed with 4% paraformaldehyde and permeabilized with 0.01% triton-x-100 before staining with antibodies against nsP4 (green), capsid (red), or DAPI (blue). Images are representative of one donor. Images were captured with the Agilent Cytation10 Confocal Imager. Scale bars = 100 µM. **(G-H)** Quantification of nsP4 positive and capsid positive macrophages detected via immunofluorescence, determined via automated counting of fluorescently positive objects. Percent infection was determined via the quotient of number of nsP4 positive or capsid positive cells divided by the number of DAPI positive cells. Data represented as means ± SEM (n= 3 donors). Statistical analysis was performed via Welch’s two-tailed T-test. Black asterisks represent P-values, where ns=non-significant, *=P<0.05, and **=P<,0.01, ***=P<0.001,

We hypothesized that differences in replication of CHIKV in GM-Mϕ and M-Mϕ might be attributed to differences in viral binding or internalization. To address this question, we performed a synchronized infection of CHIKV in GM-Mϕ and M-Mϕ to assess viral binding and entry via detection of viral nucleic acids by RT-qPCR, as previously described [48]. Briefly, macrophages were infected with a high dose of CHIKV 181/25 (MOI 5) for 2 hours at 4°C to allow binding and transferred to 37°C to facilitate subsequent viral internalization (Fig. 1C). We observed similar amounts of CHIKV RNA at the 0hr (binding) and 2hr (entry) timepoints in GM-Mϕ and M-Mϕ, suggesting that GM-Mϕ and M-Mϕ are able to bind to and internalize CHIKV virions at similar rates.

Next, we assessed expression of viral proteins in GM-Mϕ and M-Mϕ after infection with CHIKV 181/25. Immunoblotting for nonstructural protein 3 (nsP3) and the structural capsid protein revealed high expression of both proteins at 24 hours in GM-Mϕ but not in M-Mϕ (Fig 1D-1E). We confirmed this observation via immunofluorescence, where we observed significantly higher percentage of capsid and nonstructural protein 4 (nsP4) positive cells in GM-Mϕ than M-Mϕ infected macrophages (Fig 1F-1H). Notably, viral replication and expression of CHIKV viral proteins in GM-Mϕ appears to be short-lived, as replication is limited to a window between 6-24 hours. Infection of M-Mϕ could not be achieved even when supplemented with GM-CSF following initial differentiation with M-CSF (Fig S1). Interestingly, differentiation of macrophages with both GM-CSF and M-CSF resulted in higher percentage of infection than GM-CSF alone, suggesting concurrent exposure to GM-CSF and M-CSF may increase permissiveness to CHIKV.

We next evaluated whether Mayaro virus (MAYV), another arthritogenic alphavirus sharing the same receptor as CHIKV and tissue tropism [49], also preferentially infects GM-Mϕ than M-Mϕ macrophages. Consistent to our findings with CHIKV, we observed higher active replication of two MAYV strains, MAYV IQT 4235 and TRVL 4675 in GM-Mϕ macrophages at 24 hpi (Fig S2A-B). To confirm these differences in replication of MAYV IQT in GM-Mϕ and M-Mϕ macrophages, we used a reporter MAYV that expresses mCherry exclusively under the control of its subgenomic promoter and assessed infection via flow cytometry (Fig S2C). After 24 hpi, we detected significantly higher percentage of mCherry positive GM-Mϕ cells than M-Mϕ (Fig S2D-S2E). Collectively, these findings indicate that differentiation of MDMs with GM-CSF, but not M-CSF, facilitates increased susceptibility to CHIKV and MAYV infection and replication.

### Cytoskeleton and endocytic entry pathway inhibitors abrogate CHIKV infection in GM-Mϕ

Macrophages are professional phagocytes that can uptake pathogens via several endocytic pathways, including pinocytosis, micropinocytosis, and phagocytosis [50]. While the endocytic pathways by which macrophages internalize CHIKV virions are not well defined, studies using chemical inhibitors have implicated multiple endocytic mechanisms in CHIKV entry into mammalian cells [51]. To determine which endocytic pathways might be important for CHIKV infection in macrophages, we treated GM-Mϕ (since they show detectable viral protein expression) with a panel of inhibitors or vehicle control (0.1% Dimethyl Sulfoxide; DMSO) and assessed their impact on CHIKV infection (Fig 2A). Briefly, macrophages were pre-treated with inhibitors for 2 hours followed by infection with CHIKV 181/25 for 1 hour, and re-incubation with media containing inhibitors. We evaluate infection by evaluating expression of capsid protein via immunofluorescence and determined half maximal inhibitory concentration (IC50) via dose-response curve (Fig 2., orange line). We also assessed the cell viability of macrophages incubated with inhibitors via MTT (3-[4,5-dimethylthiazol-2-yl]-2,5 diphenyl tetrazolium bromide) assay and determined half maximal cytotoxic concentration (CC50), also via dose-response curve (Fig. 2, black line).

**Figure 2:**
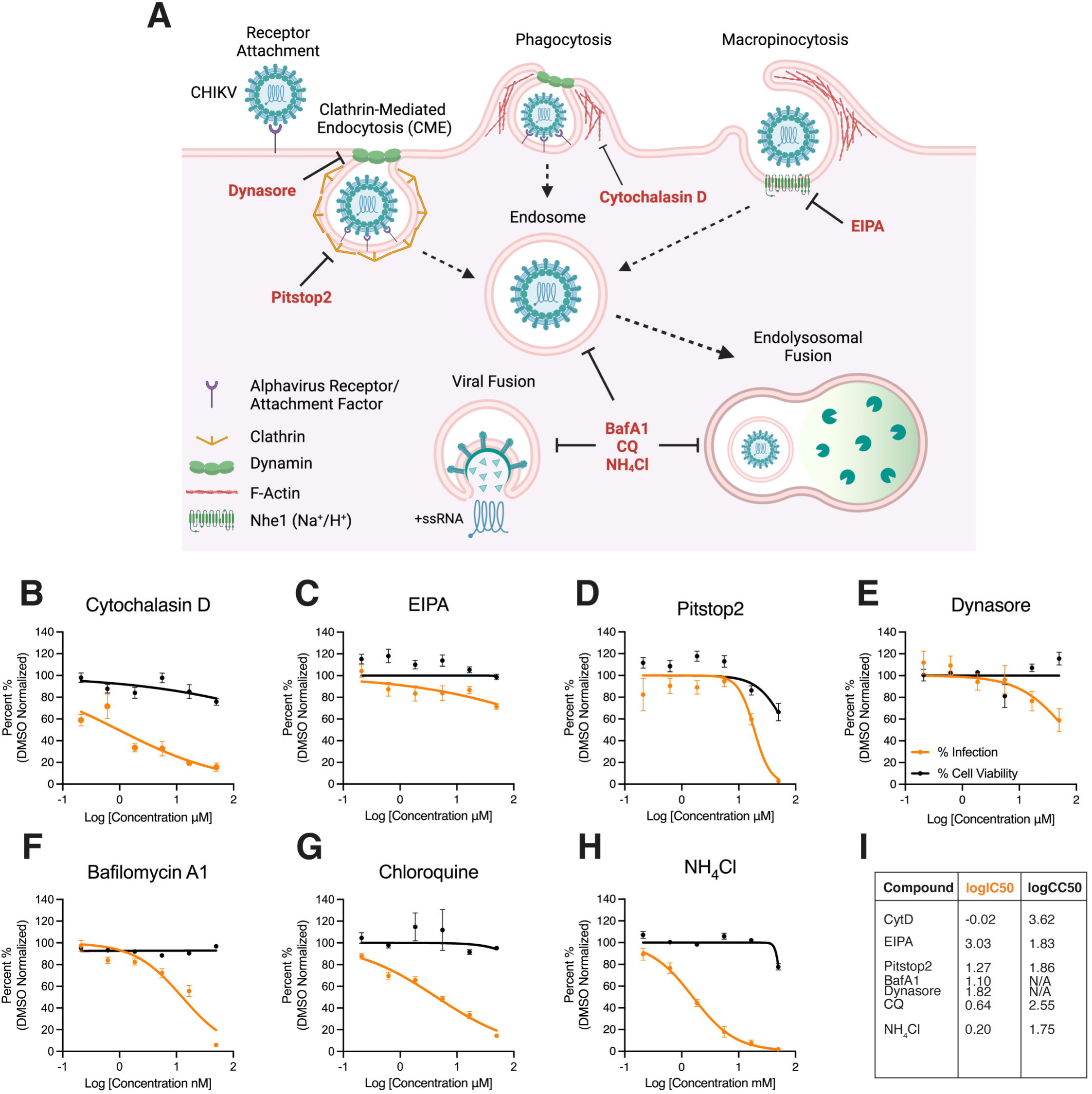
Evaluation of Cytoskeletal and Endocytic Inhibitors on CHIKV Infection in GM-Mϕ. **(A)** Schematic representation of compounds targeting CHIKV entry and internalization mechanisms in macrophages. GM-Mϕ cultured on 96-wells plates were incubated with macrophage media containing compounds inhibiting phagocytosis: **(B)** cytochalasin D, macropinocytosis: **(C)** EIPA, clathrin: **(D)** pitstop2, dynamin: **(E)** dynasore, or endocytosis/viral fusion: **(F)** bafilomycin A1, **(H)** chloroquine, **(G)** ammonium chloride, or vehicle control: DMSO for 2 hours at 37°C. Cells were then infected with CHIKV 181/25 (MOI=1.0) for 1 hour 37°C, and re-supplemented with media containing compound for a total incubation time of 24 hours. CHIKV infection was evaluated via immunofluorescence by staining for expression of CHIKV capsid protein and total cell count was determined by DAPI staining. Percent infection was determined by determining number of infected cells over total cells. Cell viability was determined via MTT assay. Data for percent (%) infection and cell viability of compounds is represented as normalized to DMSO vehicle control. **(I)** IC50 and CC50 values and corresponding dose-response curve for viral infectivity and cell viability was determined via non-linear [inhibitor] vs normalized response – variable non-linear slope analysis. Data represented as means ± SEM. Antiviral and cell viability assays were performed in technical triplicate (n=3 donors).

Phagocytosis and macropinocytosis are cellular entry processes that involve engulfment of extracellular material via cytoskeletal protrusions. Importantly, phagocytosis is a selective process that involves engagement of pathogens with phagocytic receptors while macropinocytosis is a non-specific process [52]. To assess if either of these processes are involved in CHIKV entry of macrophages, we treated GM-Mϕ with cytochalasin-D (CytD), an F-actin and cytoskeleton inhibitor which has been implicated blocking phagocytosis in macrophages [53], or with the Na+/H+ exchanger inhibitor EIPA (5-(N-ethyl-N-isopropyl)amiloride), an inhibitor of Na+/H+ exchangers (NHE) and macropinocytosis that has been shown inhibit CHIKV infection in human muscle cells [54]. We observed that treatment with CytD, but not EIPA, abrogates CHIKV infection in GM-Mϕ (Fig 2B-2C), indicating that macrophage uptake of CHIKV is a selective process.

Next, we assessed the role of clathrin-mediated endocytosis (CME) in CHIKV infection of macrophages, a process that has been heavily implicated in receptor-mediated entry of CHIKV and other alphaviruses [55]. Treatment with the clathrin inhibitor Pitstop2 abrogated CHIKV infection in GM-Mϕ, though this also led to an appreciable decrease in cell viability (Fig 2D). Dynamin is a GTPase that facilitates membrane fission in several receptor-dependent endocytic pathways, including CME and phagocytosis [56, 57]. We observed that treatment with the dynamin inhibitor dynasore abrogates CHIKV infection at the higher concentrations tested (Fig. 2E), suggesting a role for dynamin-dependent internalization of CHIKV in GM-Mϕ.

Following endocytosis, a low pH environment facilitates viral fusion with the endosomal membrane, a process essential for release of the viral genome and viral replication [58]. To confirm that CHIKV internalization in macrophages occurs via endocytosis, we treated macrophages with the V-ATPase inhibitor, bafilomycin A1, or chloroquine and ammonium chloride (NH_4_Cl); both of which are weak bases known to disrupt acidification of endosomes [58]. We observed that treatment with bafilomycin A1, chloroquine, and NH_4_Cl strongly inhibited CHIKV infection in GM-Mϕ with minimal effects on cell viability at the tested concentrations (Fig 2F-2H). Collectively, these results indicate that direct uptake of CHIKV in GM-Mϕ is a specific endocytic process likely involving multiple internalization mechanisms via dynamin- and clathrin-dependent pathways, as reflected in corresponding IC50 and CC50 values (Fig 2I). However, whether mechanisms of direct uptake of CHIKV are shared between GM-Mϕ and M-Mϕ remain elusive.

### Blocking MXRA8 and MARCO with monoclonal antibodies does not inhibit CHIKV infection in GM-Mϕ

Canonical CHIKV entry in mammalian cells involves engagement of viral glycoproteins with cell surface proteins to induce receptor-mediated endocytosis [49]. CHIKV glycoproteins have been implicated in viral entry and virion production in THP-1 macrophages and MDMs [59]. We hypothesized that GM-CSF and M-CSF might induce differential expression of surface proteins involved in alphavirus entry. To address this question, we assessed expression of a select panel of putative alphavirus receptors and entry factors in uninfected GM-Mϕ, M-Mϕ, and undifferentiated monocytes via flow cytometry [60–62]. We observed upregulation of MARCO (macrophage receptor with collagenous structure), AXL receptor tyrosine kinase, and TIM-1 (T cell immunoglobulin and mucin domain 1) upon differentiation with either GM-CSF or M-CSF but not MXRA8 (Matrix remodeling-associated protein 8) (Fig. S3A-S3B), indicating that differentiation of macrophages with GM-CSF and M-CSF selectively upregulates surface proteins implicated in alphavirus entry.

The use of MARCO as an entry receptor for alphaviruses has been shown to be species-specific, with human MARCO being dispensable for CHIKV infection [61]. In contrast, MXRA8 is well described as an entry receptor for CHIKV in human cells [62]. We tested whether blocking these receptors with monoclonal antibodies (mAbs) could inhibit CHIKV infection in GM-Mϕ, since they show detectable viral replication and viral protein expression. GM-Mϕ were incubated with various concentrations of mAbs against MARCO and MXRA8 (up to 10 μg) for 2 hours, followed by infection with CHIKV 181/25 for 1 hour, and re-incubation with media containing mAb for 24 hpi. We found that neutralization of MARCO or MXRA8 did not impact expression of CHIKV Capsid protein compared to IgG controls as detected via immunofluorescence (Fig. S3C-S3D), indicating that entry of CHIKV in MDMs is MXRA8- and MARCO- independent, though receptor-mediated entry in macrophages via other receptors cannot be ruled out.

### CHIKV induces M1-skewing of macrophage phenotype in GM-Mϕ and M-Mϕ

Differentiation of MDMs with GM-CSF or M-CSF has been shown to induce differential expression of M1- and M2-associated macrophage markers, respectively [63]. To confirm these phenotypic differences, we used spectral flow cytometry to assess surface expression of macrophage lineage, M1 pro-inflammatory, and M2 anti-inflammatory markers following differentiation.

We assessed expression of monocytic lineage markers CD11b and CD14 (Fig 3A). Expression of CD11b was upregulated in both GM-Mϕ and M-Mϕ compared to undifferentiated monocytes. In contrast, CD14 expression was higher in M-Mϕ, consistent with reported GM-CSF mediated downregulation of this receptor [64]. CD14 functions as a co-receptor for TLRs that enhances sensing of microbial and viral ligands [65], suggesting heightened innate immune sensing capacity of M-Mϕ. M1-associated markers involved in antigen presentation (CD80, CD86) were upregulated in GM-Mϕ and M-Mϕ compared to undifferentiated monocytes, whereas HLA-DR was selectively enhanced in M-Mϕ (Fig 3B). The M2-associated marker CD163 (scavenger receptor) was upregulated in both macrophage subtypes, however M-Mϕ showed higher expression post differentiation (Fig 3C). Finally, the C-type lectin receptors CD206 and CD209, which mediate ligand binding and antigen uptake via endocytosis in myeloid cells [66], were both upregulated in GM-Mϕ and M-Mϕ after differentiation.

**Figure 3:**
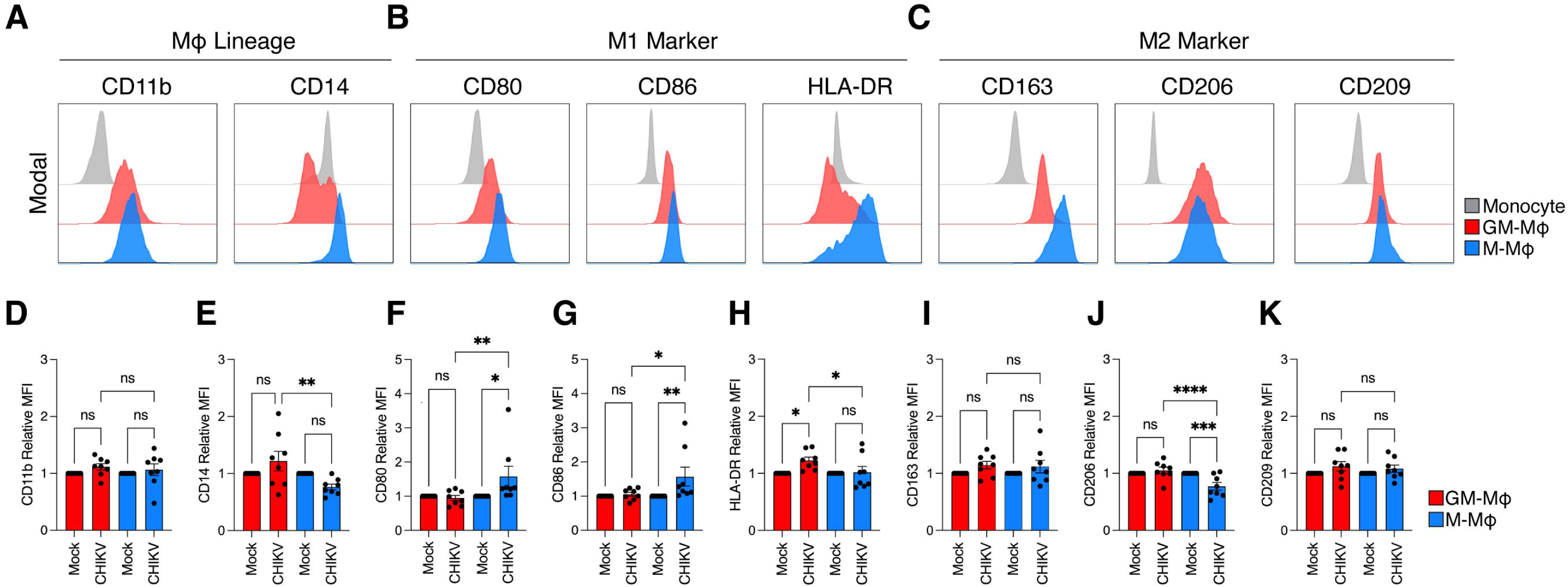
Expression of macrophage surface proteins in response to CHIKV infection in GM-M⏀ and M-MO⏀. GM-M⏀, M-M⏀, and undifferentiated monocytes were surface stained with a panel of antibodies assessing the expression of **(A)** lineage (CD11b and CD14), **(B)** M1 (CD80, CD86, and HLA-DR), and **(C)** M2 (CD163, CD206, and CD209) markers. Histograms are representative of one single donor, gated on Singlet^+^/LIVE-DEAD-Blue^-^. **(D-K)** Macrophages were infected with CHIKV 181/25 (MOI 1) or mock infected for 24 hours. Cells were stained with indicated markers. Quantification of each marker was determined by normalizing median fluorescence intensity (MFI) of live, CHIKV-infected cells over mock infected cells (n=8 donors). Data represented as means ± SEM and individual donor values. Statistical analysis was performed via one-way ANOVA with Geisser-Greenhouse correction and Fisher’s LSD test. Black asterisks represent P-values, where ns=non-significant, *=P<0.05, and **=P<0.01, ***=P<0.001, ****=P<0.0001.

We next examined whether CHIKV drives M1 or M2 skewing in GM-Mϕ and M-Mϕ by assessing relative mean fluorescence intensity (MFI) of macrophage markers after infection. CD11b expression remained unchanged in both macrophage subtypes compared to their respective mock contros (Fig 3D). Despite baseline differences, we CD14 expression was increased in GM-Mϕ after infection relative to infected to M-Mϕ, suggesting enhanced innate immune sensing in GM-Mϕ during CHIKV infection (Fig 3E). CHIKV infection induced increased expression of the M1-associated markers CD80 (Fig 3F) and CD86 (Fig 3G) in M-Mϕ but not GM-Mϕ, compared to their respective mock-treated controls. Notably, we observed significantly higher upregulation of HLA-DR in infected GM-Mϕ than M-Mϕ (Fig 3H).

The expression of M2-associated marker CD163 was unchanged in either macrophage subtype after infection (Fig 3I). Interestingly, CHIKV infection induced significantly lower expression of CD206 in M-Mϕ but not GM-Mϕ, suggesting increased receptor turnover and endocytosis during infection (Fig 3J) [67]. Finally, we detected no difference in expression of CD209 in either GM-Mϕ or M-Mϕ after infection compared to respective mock controls (Fig 3K). Collectively, these data demonstrate that CHIKV triggers a predominately M1-like activation state across both GM-Mϕ and M-Mϕ while differentially shaping phenotypic marker expression between macrophage subtypes.

### CHIKV induces a more potent type I interferon response in M-Mϕ than GM-Mϕ

CHIKV has been previously shown to induce high production of pro-inflammatory cytokines and chemokines in macrophages [26]. To investigate whether CHIKV incudes differential production of secreted cytokines and chemokines in GM-Mϕ and M-Mϕ, we evaluated the cytokine and chemokine profile of supernatants of macrophages infected with CHIKV 181/25 or UV-inactivated CHIKV 181/25 (UV-CHIKV) via multiplex enzyme-linked immunosorbent assay (ELISA). We utilized a multiplex ELISA panel consisting of pro-inflammatory and immunoregulatory cytokines and chemokines (IFNα, IL-4, IL-6, IL-10, IL-12p40, RANTES, and TNFα) known to be elevated in the serum during CHIKV infection [68]. We observed significantly higher levels of IFNα, IP10 and RANTES in M-Mϕ than GM-Mϕ after CHIKV infection (Fig 4A-4C). Notably, elevated levels of these cytokines/chemokines are only present in macrophages infected with CHIKV, but not in those exposed to UV-inactivated CHIKV, indicating that viral replication is required for their production. M-Mϕ secrete higher constitutive levels of the anti-inflammatory cytokine IL-10, and this elevated level was also maintained after infection (Fig 4D). No differences in TNFα or IL-6 production were observed between the two macrophage subtypes upon CHIKV infection (Fig 4E-4F). Collectively, these data show that differentiation of macrophages with GM-CSF and M-CSF elicits key differences in the production of antiviral cytokines and chemokines in response to CHIKV infection, possibly facilitating conditions for replication in GM-Mϕ but not M-Mϕ.

**Figure 4:**
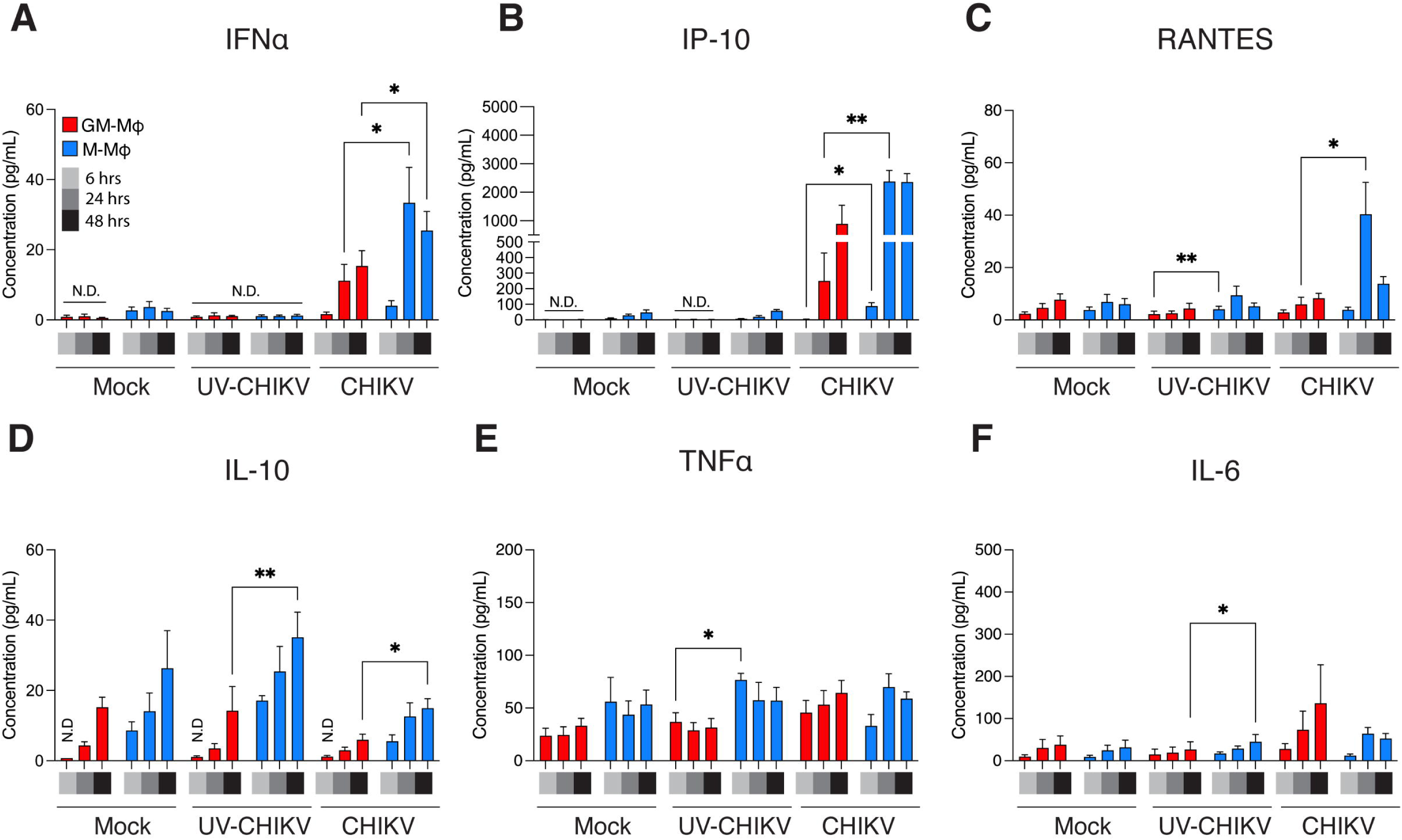
Cytokine and chemokine secretion in supernatant of primary human macrophages following CHIKV infection. Monocyte-derived macrophages were differentiated with either GM-CSF (GM-M⏀, red bars) or M-CSF (M-M⏀, blue bars) for 6 days and infected with either mock, UV-inactivated CHIKV 181/25, or CHIKV 181/25 at a multiplicity of infection (MOI) of 1.0 for 6, 24, and 48 hours. Multiplex ELISA was performed to evaluate secretion of select cytokines and chemokines: **(A)** IFN⍺, **(B)** IP-10/CXCL10, **(C)** RANTES/CCL5, **(D)** IL-10, **(E)** TNF⍺, and **(F)** IL-6. Non-detected (N.D.) indicates values below limit of detection as determined via Belysa analysis software. Data represented as means ± SEM where n=6 donors (mock and CHIKV conditions) or n=3 (UV-CHIKV condition). Statistical analysis was performed via two-way ANOVA with Geisser-Greenhouse correction and Fisher’s LSD test. Black asterisks represent P-value statistical significance, where ns=non-significant, *=P<0.05, and **=P<0.01.

To identify signaling pathways contributing to the production of pro-inflammatory cytokines and chemokines in GM-Mϕ and M-Mϕ following CHIKV infection, macrophages were stimulated with resiquimod (R848) or polyinosinic:polycytidylic acid (poly(I:C)) to selectively activate TLR7/8 or TLR3 and cytosolic RIG-I-like receptors (RLRs) sensing and signaling, respectively. Poly(I:C) treatment induced significantly higher production of IFNα and IP10 in M-Mϕ compared to GM-Mϕ, mirroring the type I IFN response observed following CHIKV infection (Fig S4A-S4B). In contrast, R848 stimulation induced robust production of RANTES, IL-10, TNFα, and IL-6 in both GM-Mϕ and M-Mϕ (Fig S4C-S4F). Notably, R848-treated GM-Mϕ produced higher levels of RANTES and TNFα, whereas R848-treated M-Mϕ exhibited higher IL-10 production. These data suggest that type I IFN production in CHIKV-infected macrophages is predominantly mediated through RLR and TLR3 signaling, with enhanced responses in M-CSF differentiated macrophages.

### CHIKV induces transcriptional activation of RNA-dependent PRR signaling cascades in M-Mϕ

CHIKV is sensed by endosomal and cytosolic pattern recognition receptors (PRRs), primarily TLR3, TLR7 and RLRs, which coordinate downstream signaling pathways in response to the diverse pathogen associated molecular patterns (PAMPs) present within the CHIKV virion [69]. To determine which innate immune sensing pathways are activated in macrophages by CHIKV, we assessed the expression of genes involved in innate immune sensing and type I IFN induction. First, we assessed expression of viral RNA sensors after infection (Fig 5A). We observed elevated gene expression of the endosomal receptors *TLR3*, *TLR7*, and *TLR8* in M-Mϕ at 24 hpi. Interestingly, we detected higher expression of the cytosolic RNA sensor *RIG-I* in GM-Mϕ compared to M-Mϕ at 24 hpi, coinciding with the elevated viral replication observed at this timepoint. We did not observe differences in CHIKV-induced *MDA5* expression between GM-Mϕ and M-Mϕ. Next, we assessed gene expression of downstream innate immune sensing adaptor proteins (Fig 5B). We observed higher expression of *TRIF*, *MYD88*, *TRAF3*, and *TRAF6* in M-Mϕ after infection. Notably, we did not observe differences in the RLR adaptor *MAVS* between GM-Mϕ and M-Mϕ. Finally, we assessed expression of innate immune transcription factors and type I IFN genes after infection (Fig 5C-5D). We detected higher expression of *IRF3* and *RELA* in M-Mϕ after at 24 hpi, while we observed no difference in *IRF7* expression between CHIKV-infected GM-Mϕ and M-Mϕ. In line with the cytokine data from infected macrophages, we observe higher expression of *IFNα* at 24 hpi and *IFNβ* at 6 hpi and 24 hpi n M-Mϕ, confirming a more robust type I IFN response in M-Mϕ than in GM-Mϕ. Collectively, these data also indicate that, compared to M-CSF, GM-CSF macrophage differentiation is associated with a reduced pro-inflammatory transcriptional response during CHIKV infection.

**Figure 5:**
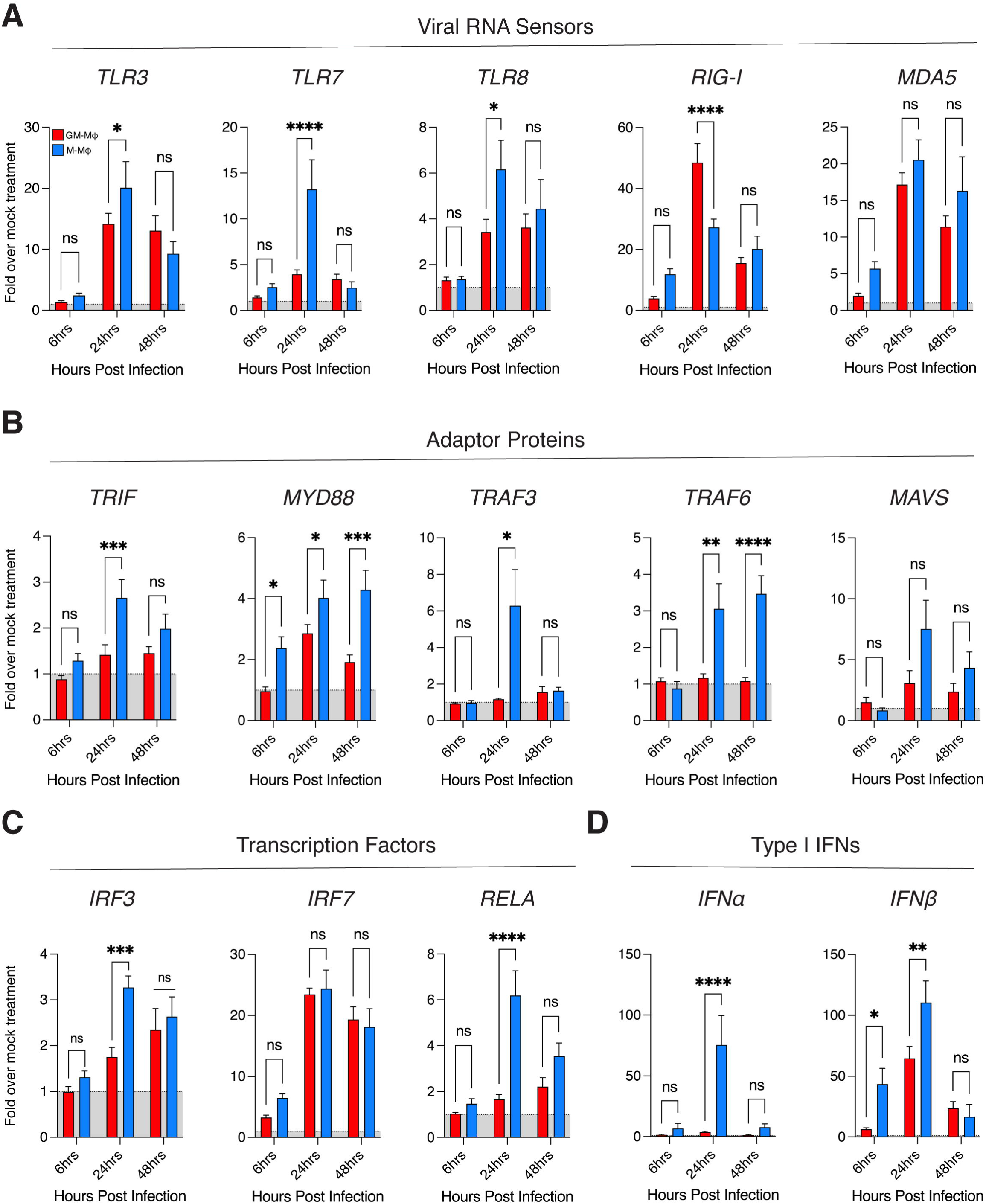
Transcriptional activation of viral RNA sensing pathways in response to CHIKV infection in GM-M⏀ and M-MO⏀. Monocyte-derived macrophages were differentiated with either GM-CSF (GM-M⏀, red bars) or M-CSF (M-M⏀, blue bars) for 6 days and with CHIKV 181/25 (MOI=1.0) for 6, 24, and 48 hours. RT-qPCR was performed to assess relative expression of the following genes: **(A)** *TLR3, TLR7, TLR8, RIG-I, MDA5* **(B)** *TRIF, MYD88, TRAF3, TRAF6, MAVS* **(C)** *IRF3, IRF7, RELA* **(D)** *IFN⍺*, and *IFNβ*. Relative expression indicates fold expression of indicated gene expression levels normalized to *RPS11* and respective mock treatment via ΔΔCT method. Data represented as means ± SEM (n= 3 donors). Statistical analysis was performed via two-way ANOVA with Geisser-Greenhouse correction and Fisher’s LSD test. Black asterisks represent P-value statistical significance, where ns=non-significant, *=P<0.05, **=P<0.01, ***=P<0.001, and ****=P<0.0001.

### CHIKV dampens interferon stimulated gene (ISG) expression in GM-Mϕ but not M-Mϕ

CHIKV viral proteins are known to inhibit type I IFN immune responses in human cells [70–72]. Since we observe higher replication and expression of CHIKV viral proteins in GM-Mϕ, we hypothesized that this might contribute to the reduced production of type I IFN responses observed in GM-Mϕ after CHIKV infection. To address this question, we infected macrophages with CHIKV followed by treatment with 1000 units of recombinant human IFNα, after which expression of the interferon stimulated genes (ISGs) *ISG15* and *MX1*, both of which are mediators of antiviral immunity and upregulated during CHIKV infection [47, 73, 74], was assessed via RT-qPCR. Treatment with IFNα alone stimulated similar robust expression of *ISG15* and *MX1* in both GM-Mϕ and M-Mϕ (Fig 6A-6B). However, infection with CHIKV prior to IFN treatment significantly dampened *ISG15* and *MX1* expression relative to respective IFNα alone treatment in GM-Mϕ but not M-Mϕ (Fig 6C-6D). These data indicate that active CHIKV replication in GM-Mϕ suppresses type I IFN signaling, whereas type I IFN signaling in M-Mϕ remains intact following infection.

**Figure 6:**
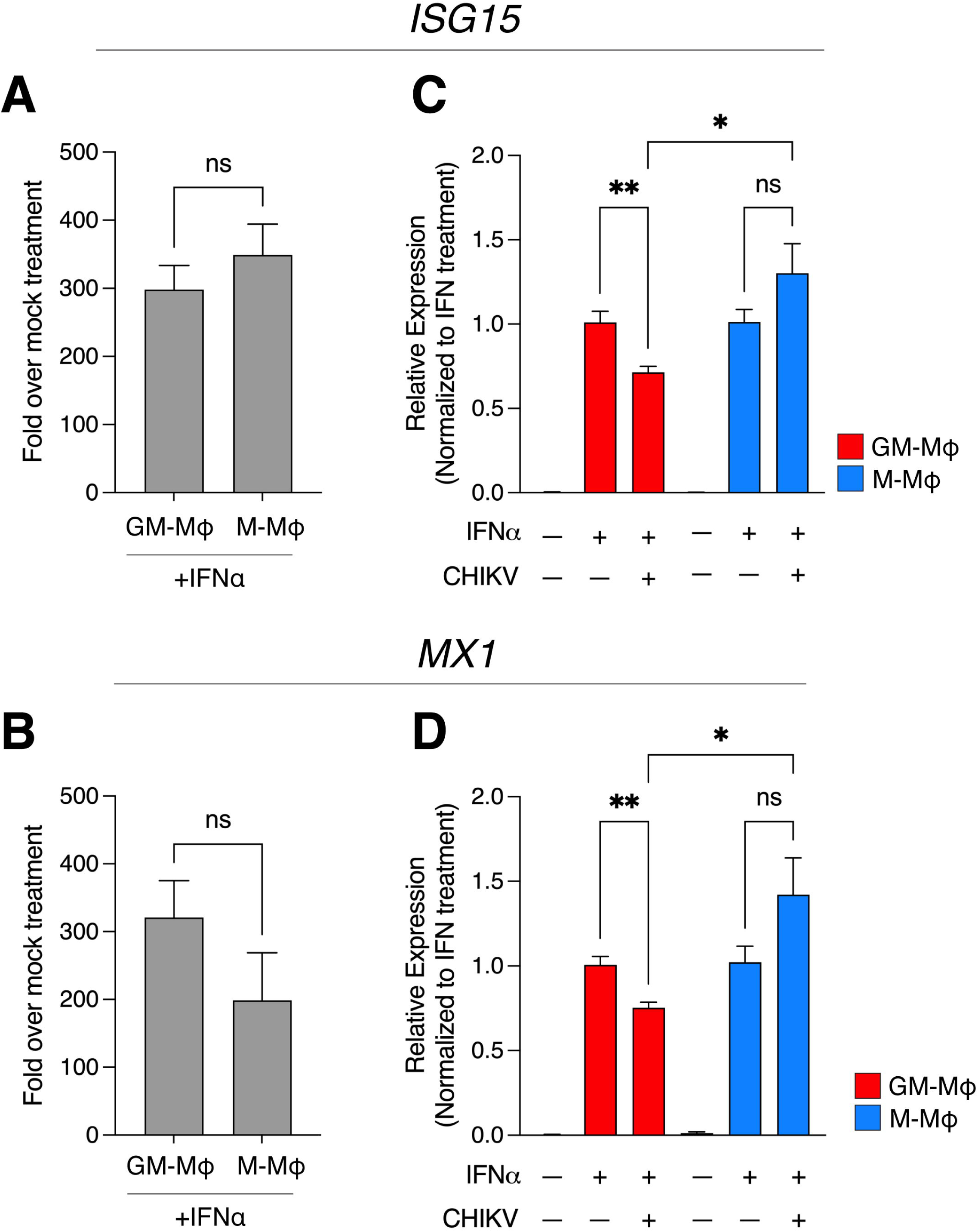
CHIKV selectively abrogates IFN⍺-induced *ISG15* and *MX1* expression in GM-M⏀ but not M-M⏀. Macrophages were treated with recombinant human IFN⍺ (1000 units/well) or vehicle control (distilled water) for 24 hours, after which cells were lysed in Trizol for RNA isolation. **(A)** *ISG15* and **(B)** *MX1* mRNA expression was evaluated via RT-qPCR analysis, where relative expression indicates fold expression of indicated gene expression levels normalized to *RPS11* and respective vehicle control treatment via ΔΔCT method. Data represented as means ± SEM (n= 3 donors). To assess effect of CHIKV-mediated inhibition of IFN⍺-induced **(C)** *ISG15* and **(D)** *MX1* expression, GM-M⏀ (red bars) and M-M⏀ (blue bars) were infected with CHIKV 181/25 (MOI=1.0) for 6 hours followed by subsequent treatment with IFN⍺ (1000 units/well) for an additional 18 hours at 37°C. After 24 hours total incubation, macrophages were lysed in Trizol for RNA isolation and RT-qPCR analysis. Data is represented as relative gene expression of ISG15 and MX1 levels normalized to *RPS11* and respective IFN⍺ alone treatment via ΔΔCT method. Data represented as means ± SEM (n= 3 donors). Statistical analysis was performed via one-way ANOVA with Geisser-Greenhouse correction and Fisher’s LSD test. Black asterisks represent P-value statistical significance, where ns=non-significant, *=P<0.05, and **=P<0.01.

## DISCUSSION

GM-CSF and M-CSF are cytokines that act as major drivers of macrophage differentiation. Using an *ex vivo* model of macrophage differentiation, we demonstrate that the alphaviruses CHIKV and MAYV establish infection and effectively replicate in GM-CSF (GM-Mϕ) but not M-CSF (M-Mϕ) differentiated MDMs. Importantly, viral entry kinetics were comparable between GM-Mϕ and M-Mϕ, indicating that resistance to CHIKV in M-Mϕ occurs post-entry. This restriction is associated with higher activation of innate immune sensing pathways in M-Mϕ that trigger type I IFN responses and establish an antiviral state. In contrast, GM-Mϕ mount a reduced antiviral response, permitting transient viral replication and expression of viral proteins. Consistent with this finding, CHIKV infection selectively antagonizes IFN-α–induced *ISG15* and *MX1* gene expression in GM-Mϕ, but not in M-Mϕ, highlighting differentiation-dependent differences in innate immune control of infection in human macrophages.

The contribution of macrophages as reservoirs for persistent alphavirus infection represents a major area of investigation in defining the immunopathogenic mechanisms underlying CHIKVD [75]. In this study, we demonstrate that GM-CSF facilitates conditions in MDMs that increases susceptibility to active replication and antigen production of CHIKV and MAYV. *In vivo* data in mice indicates that CHIKV infection induces high production of GM-CSF and IFNγ by activated CD4^+^ T cells, maintaining a pro-inflammatory environment that enables GMCSF-differentiation of infiltrating macrophages in joint tissue and driving pathogenesis [35]. Thus, crosstalk between macrophages and GMCSF-producing cells may create a pro-inflammatory cytokine milieu that facilitates macrophage alphavirus infection and persistence. Our data also suggests that GM-CSF may have a dominant role in macrophage programming, as we demonstrate that monocytes initially differentiated in the presence of GM-CSF remain highly susceptible to CHIKV infection following subsequent or concurrent exposure to M-CSF. However, because GM-CSF, M-CSF, and other macrophage differentiation factors are likely simultaneously present within inflamed tissue during CHIKV infection, *in vivo* studies must be performed to fully understand macrophage differentiation and the contribution of other cell types and activation signals during infection. Additionally, the role of GM-CSF and M-CSF in the context of human tissue-resident macrophages during CHIKV infection remains unexplored.

Understanding CHIKV entry into macrophages is essential for identifying host factors that enable them to serve as viral reservoirs. In this study, we investigated macrophage entry mechanisms involved in CHIKV infection using chemical inhibitors. Treatment with inhibitors for phagocytosis, dynamin- and clathrin- dependent entry markedly abrogated CHIKV infection in GM-Mϕ, whereas treatment for non-specific uptake via macropinocytosis did not reduce infection at the range of concentration tested. Additionally, inhibition of viral fusion strongly suppressed infection, collectively implicating endocytic entry as a critical requirement for productive CHIKV infection in macrophages. Importantly, bafilomycin A1, chloroquine, and NH_4_Cl are lysosomotropic compounds that broadly disrupt pH-dependent endocytic and lysosomal processes not explored in this study, including autophagy, which has been shown to exert both anti- and pro-viral effects [76]. Moreover, differential regulation of autophagy and lysosomal trafficking by GM-CSF and M-CSF may contribute to the distinct susceptibilities to CHIKV infection observed between the two macrophage subtypes [77].

Macrophages exhibit a high degree of plasticity, enabling them to adopt pro- or anti-inflammatory effector functions in a context-dependent manner, which is reflected in the diverse repertoire of surface markers expressed across macrophage subtypes [78, 79]. Although GM-CSF and M-CSF drive distinct differentiation programs that bias macrophages toward M1-like or M2-like states, respectively, we find that CHIKV induces a predominantly M1-associated activation profile in both macrophage subtypes, as mainly evidenced by upregulation of co-stimulatory and antigen-presentation markers in both in both GM-Mϕ and M-Mϕ. We also observe significantly lower surface expression of the M2-associated marker CD206 in CHIKV-infected M-Mϕ, further suggesting M1-skewing as turnover of this receptor has been linked to increased antigen presentation [67]. Notably, internalization of CD206 has been associated with endocytosis of pathogens, including viruses [80], though a mechanism for receptor-mediated CHIKV entry in macrophages via CD206 was not explored in this study. Additionally, blocking alphavirus receptors with mAbs against MXRA8 or MARCO did not inhibit CHIKV infection in GM-Mϕ, suggesting other receptors might be important for CHIKV entry in macrophages.

M1-activated macrophages have been described to be susceptible to infection by multiple RNA viruses, including dengue virus (DENV), SARS-CoV-2, and Mumps virus (MuV) [81–83]. Despite exhibiting a M1-activated state during CHIKV infection, M-Mϕ remain resistant to productive CHIKV replication, underscoring the complexity of macrophage differentiation beyond the classical GM-CSF/M1 and M-CSF/M2 paradigm. Emerging evidence also suggests that human macrophage activation may vary during CHIKVD progression [84], however, the relative contributions of GM-CSF and M-CSF to macrophage phenotype at distinct disease stages have yet to be defined.

Skewing of macrophages towards M1-activation during CHIKV infection is likely driven by differential priming of innate immune sensing pathways by GM-CSF and M-CSF, which has been previously shown in macrophages during West Nile Virus (WNV) infection [85]. Previous studies have shown that M-Mϕ exhibit higher basal and interferon-induced levels of TLR3 and TLR7 than GM-Mϕ [86–88]. Here, we demonstrate that, during CHIKV infection, gene expression of endosomal receptors *TLR3*, *TLR7*, and *TLR8* are more highly upregulated in M-Mϕ than GM-Mϕ. Notably, cytosolic RLR *RIG-I* is more strongly upregulated in actively replicating GM-Mϕ than M-Mϕ and *MDA5* is similarly activated in both subtypes, suggesting that RLR activation may not be the primary driver of the heightened type I IFN response in M-Mϕ. Indeed, we show that stimulation with poly(I:C) but not R848 triggers production of IFNα and IP10 in both GM-Mϕ and M-Mϕ, indicating that antiviral activation of macrophages is likely driven via TLR3 signaling. This data is consistent with other studies investigating innate immune responses to CHIKV in primary macrophages, which also show antiviral activation of macrophages via TLR3-dependent production of IL-27 [24, 26]. The loss of TLR3 has been shown to increase CHIKV viremia in a TLR3-knockout mouse model and polymorphisms of TLR3 in CHIKV-infected human PBMCs has been linked with increased disease severity, further emphasizing the importance of TLR3 in controlling CHIKV replication [89].

Surprisingly, CHIKV infection resulted in a muted pro-inflammatory cytokine and chemokine response in both GM-Mϕ and M-Mϕ, with the exception of RANTES/CCL5, which was selectively elevated in infected M-Mϕ. Similarly, stimulation of GM-Mϕ and M-Mϕ with poly(I:C) induced elevated IFNα and IP10 production alongside a suppressed pro-inflammatory cytokine and chemokine response, mirroring the profile observed during infection. Interestingly, the lack of pro-inflammatory cytokine induction following TLR3 activation appears to be cell type and species-specific. Indeed, poly(I:C) has been previously reported to induce IP10 but not TNFα or IL-6 in human monocyte-derived macrophages and dendritic cells, whereas human synovial fibroblasts and murine dendritic cells produce robust TNFα and IL-6 responses [90]. Additionally, the significantly higher production of IFNα and IP10 in M-Mϕ compared to GM-Mϕ during infection and poly(I:C) treatment suggests that these outcomes are dictated primarily by intrinsic features of macrophage differentiation rather than specific immune modulation by CHIKV. Indeed, despite its described roles in pro-inflammatory activity, GM-CSF has been shown to dampen certain TLR expression and signaling compared to non-treated monocytes [91], highlighting the complexity of cytokine-driven regulation of innate immune responses. In addition, differential expression of negative regulators of innate signaling, such as suppressor of cytokine signaling (SOCS) proteins induced by GM-CSF or M-CSF, may contribute to these effects by potentially attenuating TLR, NF-κB, and JAK/STAT pathway responsiveness [92–96]. While speculative, this model is consistent with prior reports describing viral exploitation of myeloid differentiation programs via SOCS to facilitate replication and immune evasion [97, 98]. Additional transcriptomic and proteomic analyses will be required to define the molecular programs underlying GM-CSF-driven permissiveness and M-CSF-associated antiviral restriction to CHIKV.

Our findings suggest that macrophage differentiation state may influence acute viral control and inflammatory responses to infection. Collectively, our findings identify GM-CSF-driven macrophage differentiation as a determinant of CHIKV permissiveness with potential contributions to viral persistence. By enabling transient viral replication and blunting type I IFN signaling, GM-CSF primed macrophages may contribute to dissemination of CHIKV to peripheral organs and tissue, potentially contributing to disease pathophysiology. How GM-CSF and M-CSF shape inflammatory responses in the development of chronic CHIKVD is a critical future direction necessary to understand clinical relevance beyond the acute stage of disease. Together, our results highlight macrophage differentiation state as a critical factor in CHIKV pathogenesis and suggest that therapeutic modulation of GM-CSF and M-CSF signaling may represent a strategy to limit infection while minimizing inflammatory damage in CHIKV disease.

## MATERIALS AND METHODS

### Differentiation of primary macrophages

Monocyte-derived macrophages (MDMs) were generated from buffy coats of healthy human donors obtained from the New York Blood Center. Buffy coats underwent Ficoll-Hypaque centrifugation followed by isolation of CD14+ cells using a MACS CD14 isolation kit (Milteny Biotec) according to manufacturer’s instructions. CD14+ monocytes were cultured in RPMI 1640 medium supplemented with 10% hyclone fetal bovine serum, 100 U mL penicillin/streptomycin, 10 mM HEPES, and 1 mM sodium pyruvate (NaPyr). Monocytes were differentiated into GM-CSF derived (GM-Mϕ) and M-CSF derived (M-Mϕ) macrophages via incubation for six days in media containing 100 U/mL human recombinant granulocyte-macrophage colony stimulating factor (GM-CSF) (Peprotech) or 100 U/mL human recombinant macrophage colony stimulating factor (M-CSF) (STEMCELL #78057), respectively.

### Cell Lines

U2OS human bone osteosarcoma epithelial cells were gifted by Dr. Carolyn Coyne and were cultured in Dulbecco’s Modified Essential Medium (DMEM) supplemented with 10% fetal bovine serum, 100 U mL L-glutamine, and 100 U mL penicillin/streptomycin. HFF-1 fibroblast cells were cultured in Dulbecco’s Modified Essential Medium (DMEM) supplemented with 20% fetal bovine serum.

### Biosafety

Experiments involving CHIKV AF15561 were performed in a biosafety level 3 (BSL-3) laboratory at the Icahn School of Medicine at Mount Sinai.

### Viruses

Viruses from Infectious clones encoding for Chikungunya virus (CHIKV AF15561) and Mayaro virus (MAYV-TRVL 4675, MAYV-IQT 4235, and MAYV-IQT 4235-mCherry) were generated from infectious clones and rescued as previously described [47]. Infectious clones of CHIKV AF15561 and MAYV-IQT 4235 were kindly provided by the World Reference Center for Emerging Viruses and Arboviruses (WRCEVA). The MAYV-TRVL 4675 infectious clone was a generous gift from Dr. Andres Merits. Chikungunya 181/25 (CHIKV 181/25) strain is an attenuated vaccine strain that was originally isolated from a patient in Thailand. An aliquot of CHIKV 181/25 which was provided by Dr. St. Patrick Reid at the University of Nebraska and has been passaged twice in Vero cells and once in BHK cells. Supernatants were centrifuged at 1500 RPM for 10 minutes and stored at - 80°C in single-use 0.5 mL aliquots. UV inactivation of CHIKV and MAYV was performed as previously described[70].

### Infection and stimulation of MDMs

After differentiation, donor-matched macrophages were mock-infected with blank RPMI media or with virus diluted in blank RPMI media to achieve desired multiplicity of infection (MOI). Cells were infected for one hour at 37°C, rocking the plate to distribute the inoculum across the wells every 15 minutes. After the hour infection, the inoculum was removed and cells were supplemented with appropriate fresh media and incubated at 37°C for up to 48 hours. Stimulation of MDMs was performed via incubation of cells in macrophage media supplemented with R848 (resiquimod) (Sigma) or poly(I:C) (Sigma) for indicated timepoints.

### RNA isolation

Primary macrophages were lysed an inactivated in trizol (Thermo) and stored at -80°C until ready for RNA isolation. Samples lysed in trizol were thawed at room temperature and RNA was extracted using Zymo Direct-zol RNA Isolation kit following manufacturer’s instructions, including in-column DNase treatment. RNA was eluted in UltraPure DNase/RNase free distilled water and stored at -80°C until future use. The concentration and quality of RNA was determined via measurement on a Thermo Scientific Nanodrop One spectrophotometer at 260 nm.

### Reverse Transcription-quantitative polymerase chain reaction (RT-qPCR)

Following RNA isolation, RT-qPCR was used to quantify relative gene expression of select innate immune signaling genes using the New England BioLabs Luna Universal One-Step RT-qPCR Kit, per manufacturer’s instructions. PCR was performed on the BioRad 1000C thermal cycler on the SYBR/FAM scan mode with the following thermocycling protocol: (1) reverse transcription for 1 cycle at 55°C for 10 minutes, (2) initial denaturation for 1 cycle at 95°C for 1 minute, (3) denaturation at 95°C for 10 seconds and (4) extension at 60°C for 30 seconds for 40 cycles. A melt curve step was added to assess specificity of amplification. Relative expression of innate immune genes was determined via normalization to the housekeeping gene *RPS11* and respective mock treatment via the ΔΔCT method.

### Viral RNA quantification

Quantification of CHIKV and MAYV RNA concentration in primary macrophages was determined via a standard curve of *in vitro* transcribed (IVT) RNA, as previously described [47]. A standard curve was performed for every 96-well PCR plate spanning a concentration range of 50 ng/μL to 5E-6 ng/μL. RT-qPCR (SYBR/FAM) was then performed to determine CT values of the viral genes *E1* and *nsP1* using CHIKV and MAYV specific primers. Concentration of viral transcripts in the sample was determined via normalization to the concentration of total RNA in each sample and total RNA in each reaction. Active replication was determined by quantifying the ratio of subgenomic RNA (E1) to genomic RNA (nsP1).

### CHIKV binding and entry assay

MDMs were seeded and differentiated on 96-well plates and infected with CHIKV 181/25 (MOI 5) for 2 hours on ice to synchronize infection. The following conditions were performed on separate plates: binding assessment was performed by washing cells with ice-cold PBS 3x and lysed in trizol immediately following 2-hour synchronized infection. Assessment of viral internalization was performed by incubating cells at 37°C for 2 hours following synchronized infection and then washing with PBS 3x and lysed in trizol. Viral RNA concentration was determined via standard curve, as explained above.

### Co-treatment of MDMs

MDMs were infected with CHIKV 181/25 (MOI 1) or mock infection for 6 hours at 37°C to allow initiation of transcription and translation of viral proteins. Following incubation, cells were co-treated with MDM media containing 1000 units of recombinant human IFNα (Thermo) or vehicle control (distilled water). Cells were then incubated at 37°C for an additional 18 hours, for a total incubation period of 24 hours. Cells were then lysed in trizol and RNA was isolated as previously described. Expression of *ISG15*, *MX1*, and *RPS11* (housekeeping gene) was then measured via RT-qPCR. Type I IFN signaling antagonism was assessed via normalizing all conditions to *RPS11* and then determining fold induction of secondary treatment over the respective mock-infected IFNα treatment via the ΔΔCT method. For example, determining expression levels of *ISG15* following co-treatment of CHIKV and IFNα was determined by comparing the following conditions: CHIKV → IFNα over Mock → IFNα (primary→secondary).

### Aurora Spectral Flow Cytometry

MDMs were seeded and differentiated on 24-well plates at a cell density of 5 x 10^5^ cells/mL. At the time of collection, media was aspirated and cells were washed with ice-cold PBS and incubated with 5 mM EDTA in PBS for 20 minutes to allow for cell detachment. After incubation, cells were fully detached via gentle pipetting and transferred to 1.5 mL Eppendorf microcentrifuge tubes. Cells were spun at 1500 G for 5 minutes at 4°C. Cells were then resuspended in PBS and transferred to a 96-well plate (round bottom) to continue flow cytometry staining. Cells were stained for viability with 100 μL of LIVE/DEAD fixable Blue (1:2000, Thermo) for 15 minutes on ice. Cells were washed with FACS buffer (PBS supplemented with 2% FBS and 5 mM EDTA) and stained with Human TruStain FcX blocking solution (5 μL per 1 x 106 cells) (BioLegend) diluted in FACS buffer for 15 minutes on ice. Cells were then stained for 1 hour on ice with a cocktail of the following antibodies: CD11b-BV650 (BioLegend), CD14-APC eFluor 780 (Invitrogen), CD80-BUV737 (Invitrogen), CD86-BV480 (Invitrogen), HLA-DR-BUV805 (Invitrogen), CD163-PE (BioLegend), CD206-BV711 (BioLegend), and CD209-PE-Cy7 (BioLegend). Alternatively, for analysis of CHIKV receptor surface expression, cells were stained with a cocktail of the following antibodies: TIM-1/CD365-PE-Cy7 (BioLegend), MARCO-PE (Invitrogen), and AXL-APC (Invitrogen). Staining of MXRA8 was performed via two-step process with an unconjugated mAb for MXRA8, 2H2G12A (1:100, Medical & Biological Laboratories Co) and secondary staining with Alexafluor-488 goat anti-rabbit IgG (H+L) (1:1000, Invitrogen), washing extensively between staining steps. After incubation, cells were washed twice in FACS buffer and centrifuged at 1500 G for 5 minutes. Cells were then fixed with 100 μL of BD Biosciences perm fixation buffer (BD Biosciences) on ice for 20 minutes. Cells were then washed twice in FACS buffer and centrifuged at 1500 G for 5 minutes and resuspended in 200 μL FACS buffer for acquisition on the Cytek Aurora Spectral flow cytometer. Flow cytometry analysis was performed on FlowJo analysis software.

### Immunofluorescence

Infected MDMs were fixed with 4% paraformaldehyde (PFA) diluted in PBS for 20 minutes at room temperature. Cells were then permeabilized with 0.1% Triton-X-100 for 10 minutes. Following permeabilization, cells were washed with PBS and blocked with blocking buffer (4% BSA in PBS) for 1 hour at room temperature. Cells were then stained with primary antibodies for CHIKV capsid protein (1:1000, goat anti-mouse Millipore Sigma) and CHIKV nsp3 (1:1000, goat anti-rabbit, gift from Dr. Kenneth Stapleford) for 1 hour at room temperature in blocking buffer. Cells were then stained with the following secondary antibodies: Alexafluor-488 goat anti-rabbit IgG (H+L) (1:1000, Invitrogen) and Alexafluor-647 goat anti-mouse IgG (H+L) (1:1000, Invitrogen) for 1 hour in the dark at room temperature. Cells were then counterstained with PBS supplemented with DAPI (1:10000). Cells were washed with PBS 3x between each step in the staining procedure, except between the blocking and primary antibody staining step. Plates were imaged on the Agilent BioTek Cytation10 Confocal Imaging Reader using the imager manual mode at 10x magnification. Initial image deconvolution was performed within the Gen5 Image+ software and final review and processing was performed on ImageJ. Quantification was performed via the microplate reader and scanning feature on the Cytation10. Wells were scanned at 4x magnification and infected cells were quantified by counting Capsid+ or nsp3+ cells over the total number of cells (DAPI+) in each respective well.

### Inhibitor Treatment and Cell Viability Assay

MDMs were seeded and differentiated in duplicate 96-well plates for antiviral assay (AV) and cell viability assay (CV). Serial dilutions of compounds (1:3 dilution series) were performed in deep-well plates containing MDM media. Cells were treated with media containing compound or vehicle control (DMSO) 2 hours (AV plates) or 24 hours (CV plates) at 37°C. After 2-hour incubation, media was aspirated from AV plates and infected with 50 μL of CHIKV 181/25 (MOI 1) and placed back in the incubator for 1 hour. After infection, inoculum from AV plates was aspirated and cells were re-supplemented with media containing compound and incubated for an additional 22 hours. After 24 hours, AV plates were washed with PBS and fixed with 4% PFA and underwent immunofluorescence staining to determine percent CHIKV infectivity, as previously explained. CV plates underwent MTT-based colorimetric cell proliferation assay (Roche), per manufacturer’s instructions. Spectrophotometrical absorbance was measured with the Cytation10 microplate reader at 550 nm and a reference wavelength of 750 nm. Cell viability percentage was determined by normalizing absorbance values of compound treated cells to vehicle control treated cells. Dose-response curves of CHIKV infectivition and cell viability were determined via non-linear [inhibitor] vs normalized response – variable non-linear slope analysis. IC50 (half maximal inhibitory concentration) of viral infection inhibition was determined via best-fit concentration value of dose-response curve at fifty percent inhibition. Lysosomotropic compounds (bafilomycin A1, chloroquine, and ammonium chloride) were obtained from Thermo. Compound targeting phagocytosis (cytochalasin D) was obtained from Sigma-Adlrich. Other compounds targeting micropinocytosis (EIPA), clathrin (pitstop 2), and dynamin (dynasore) were obtained from MedChemExpress. All compounds were reconstituted in DMSO.

### Macrophage Receptor Neutralization Assay

MDMs were seeded and differentiated in 96-well plates. Serial dilution of mAbs (1:5 dilution series) were performed in deep-well 96-well plates containing MDM media. The following antibodies were used for this assay: MXRA8 (2H2, Medical & Biological Laboratories Co), MARCO (PLK1, Novus Biologicals), mouse IgG2 or IgG3 isotype control (Cell Signaling). Cells were treated with media containing antibody for 2 hours at at 37°C. After 2-hour incubation, media was aspirated from and cells were infected with 50 μL of CHIKV 181/25 (MOI 2) and placed back in the incubator for 1 hour. After infection, inoculum from plates was aspirated and cells were re-supplemented with media containing antibody and incubated for an additional 22 hours. After 24 hours, plates were washed with PBS and fixed with 4% PFA and underwent immunofluorescence staining to determine percent CHIKV infectivity, as previously explained.

### Immunoblot Analysis

Cellular protein lysates were obtained by lysing cells in RIPA buffer (Sigma Aldrich) supplemented with 1x PhosoSTOP phosphatase inhibitor (Roche) and 1x EDTA-free cOmplete Ultra protease inhibitor (Roche) for 15 minutes on ice. Protein lysates were loaded onto pre-cast 4-20% poly acrylamide SDS gels (Bio-Rad) and separated via SDS-PAGE electrophoresis. Gels were then transferred onto nitrocellulose membranes (Bio-Rad) via semi-dry transfer using Trans-blot turbo transfer system (Bio-Rad). Membranes were then transferred to a platform rocker and washed in washing buffer (1x Tris-buffered saline + 1% tween-20) (Cell Signaling) and blocked for 1 hour at room temperature in blocking buffer (5% BSA in TBS-T washing buffer) (Thermo). Membranes were then probed with primary antibodies diluted in blocking buffer overnight at 4°C. The following day, membranes were washed for 5 minutes 3x in washing buffer and probed with appropriate secondary antibody diluted in blocking buffer for 1 hour at room temperature. Membranes were then washed for 5 minutes 3x in washing buffer. Membranes were then dried and exposed using Femto or Pico chemiluminescent substrate reagent (Thermo) for 5 minutes. Detection of protein expression signal was performed using a digital developer. Primary antibodies used: anti-β-Actin (1:5000, Sigma-Aldrich), anti-CHIKV CP (1:1000, Millipore Sigma), anti-CHIKV nsP3 (1:3000, polyclonal antibody donated by Dr. Kenneth A. Stapleford), anti-MXRA8 (1:1000, Cell Signaling). Secondary antibodies used: Goat Anti-Mouse IgG (1:5000, Sigma-Aldrich), Goat Anti-Rabbit IgG (1:5000, Sigma-Aldrich).

### Multiplex ELISA

Cytokine and chemokine analysis was performed as previously described [99]. Briefly, a cytokine magnetic 9-plex panel for the Luminex platform (Millipore Millplex) was used according to manufacturer’s instructions. The cytokine panel consisted of the following analytes: IFNα, IP10, TNFα, IL-6, IL-8, IL-10, RANTES/CCL5, MIP1B, and VEGFα (note: not all cytokines were used in the final analysis). The cytokine concentrations were then determined using Belysa Immunoassay Curve Fitting Software (V1.1.0). Data analysis was performed using the Millplex Analyst software.

### Statistical analysis

Statistical analysis was performed with GraphPad PRISM software (version 10) using Welch’s two-tailed T-test for direct comparisons or ANOVA (one-way or two-way, where indicated) with Geissler-Greenhouse correction followed with LSD Fisher’s multiple comparison test to determine the significance of variance among indicated groups. P-values are indicative of group comparisons, where P<0.05(*), P<0.01(**), P<0.001(***), P<0.0001(****), and P>0.05(non-significant/ns). Prior statistical determination of sample size was not considered in this study.

## ACKNOWLEDGMENTS

We would like to thank Chris Basler and Kris White for technical support with the Cytation imager and plate reader and antiviral assays. We also thank Jeffrey Johnson and Boris Bonaventure for reagents and insightful discussion. In addition, we thank Randy Albrecht and Lokendrasingh Chauhan for their biosafety expertise and support in managing and maintaining the BSL-3 facility at Mount Sinai. Finally, we thank the Mount Sinai flow cytometry core for their support with the Aurora spectral flow cytometry.

## Supporting Information

**S1 Fig.** Temporal addition of GM-CSF or M-CSF on CHIKV infectivity in primary macrophages. Monocyte-derived macrophages were differentiated with GM-CSF, M-CSF, or both cytokines and infected with CHIKV 181/25 (MO1=1.0) for 24 hours. To assess whether susceptibility to CHIKV can change, macrophages were treated with the opposite cytokine at 0-, 3-, or 5-days post differentiation. Quantification of capsid positive macrophages was detected via immunofluorescence. Percent infection was determined via the quotient of number of capsid positive cells divided by the number of DAPI positive cells. Data represented as means ± SEM (n= 2 donors). Statistical analysis was performed via one-way ANOVA with Fisher’s LSD test. Statistical analysis was performed via one-way ANOVA with Geisser-Greenhouse correction and Fisher’s LSD test. Black asterisks represent P-value statistical significance, where ns=non-significant, *=P<0.05, **=P<0.01, ***=P<0.001, and ****=P<0.0001.

**S2 Fig.** Viral infectivity and replication of Mayaro virus in GM-M⏀ and M-M⏀. (A-B) GM-M⏀ (red bars) or M-M⏀ (blue bars) for 6 days and infected with MAYV TRVL or MAYV IQT (MOI=1.0) for 6, 24, and 48 hours. Intracellular viral replication is represented as “active replication”, which was determined by evaluating the ratio of E1 subgenomic viral RNA copies to nsP1 genomic viral RNA copies. Concentration of viral RNA was determined by RT-qPCR via standard curve. Statistical analysis was performed via two-way ANOVA with Geisser-Greenhouse correction and Fisher’s LSD test. Data represented as means ± SEM (n=4 donors). (C) Schematic representation of infectious clone encoding MAYV-IQT-mCherry reporter virus, where the mCherry protein is expressed under the control of a subgenomic promoter. (D) GM-M⏀ (top row) and M-M⏀ (bottom row) were infected with mock, UV-inactivated MAYV IQT-mcherry (MOI=1.0), or MAYV IQT-mCherry (MOI=1.0) for 24 hours and prepared for flow staining. Flow plots represent mCherry+/LIVE-DEAD-/Singlet. Flow plots are representative of one donor. (E) Quantification of mCherry+ mock, UV-inactivated MAYV IQT-mCherry, or MAYV IQT-mCherry GM-M⏀ and M-M⏀ determined via flow cytometry. Data represented as means ± SEM (n=4 donors). Statistical analysis was performed via one-way ANOVA Geisser-Greenhouse correction and Fisher’s LSD test. Black asterisks represent P-values, where ns=non-significant, *=P<0.05, and **=P<,0.01, ***=P<0.001, ****=P<0.0001.

**S3 Fig.** Evaluation of alphavirus receptor expression in GM-M⏀ and M-M⏀. GM-M⏀ (red), M-M⏀ (blue), and undifferentiated monocytes (gray) were surface stained with a panel of antibodies assessing the expression of alphavirus receptors (A) MARCO, MXRA8, AXL, and TIM-1 (CD365). Histograms are representative of a single donor. (B) Immunoblot analysis of total MXRA8 and β-Actin protein expression in un-infected U2OS, HFF-1, GM-M⏀ and M-M⏀. Data is representative a single replicate (cell lines) and single donor (macrophages). GM-M⏀ were pre-treated with media containing mAbs (up to 10 μg/mL) against (C) MXRA8 or (D) MARCO and respective IgG isotype controls. Cells were then infected with CHIKV 181/25 (MOI=2.0) for 1 hour, followed by re-supplementation with media containing antibodies for a total incubation time of 24 hours. To determine percent infection, immunofluorescence staining of CHIKV capsid protein to determine the number of infected cells and DAPI to determine total cell count. Infection data from antibody treatment is represented as relative percent infection normalized to DMSO control treatment (n=4 donors). Statistical analysis was performed via two-way ANOVA Geisser-Greenhouse correction and Fisher’s LSD test. Black asterisks represent P-values, where ns=non-significant.

**S4 Fig.** Cytokine and chemokine secretion in supernatant of GM-M⏀ and M-M⏀ following R848 or poly(I:C) stimulation. GM-M⏀ (red bars) or M-M⏀ (blue bars) for 6 days and treated with 1 μg/mL of R848 or poly(I:C) for 6, 24, and 48 hours. Multiplex ELISA was performed to evaluate secretion of select cytokines and chemokines: (A) IFN⍺, (B) IP-10/CXCL10, (C) RANTES/CCL5, (D) IL-10, (E) TNF⍺, and (F) IL-6. Non-detected (N.D.) indicates values below limit of detection as determined via Belysa analysis software. Data represented as means ± SEM where n=4 donors. Statistical analysis was performed via two-way ANOVA with Geisser-Greenhouse correction and Fisher’s LSD test. Black asterisks represent P-value statistical significance, where *=P<0.05, and **=P<0.01. Statistical comparisons resulting in non-significant P-values (>0.05) were performed but not reported.

## REFERENCES

1. de Souza WM, Ribeiro GS, de Lima STS, de Jesus R, Moreira FRR, Whittaker C, et al. Chikungunya: a decade of burden in the Americas. The Lancet Regional Health – Americas. 2024;30. doi: 10.1016/j.lana.2023.100673.

2. Grabenstein JD, Tomar AS. Global geotemporal distribution of chikungunya disease, 2011–2022. Travel Medicine and Infectious Disease. 2023;54:102603. doi: 10.1016/j.tmaid.2023.102603.

3. Yang Y-F, Qiu Y-B, Xu Q, Gao R-C, Tang T, Tian Y, et al. Mapping the global risk of chikungunya virus endemicity and autochthonous transmission following importation. Travel Medicine and Infectious Disease. 2025;67:102892. doi: 10.1016/j.tmaid.2025.102892.

4. Tee KK, Mu D, Xia X. Explosive chikungunya virus outbreak in China. International Journal of Infectious Diseases. 2025;161:108089. doi: 10.1016/j.ijid.2025.108089.

5. Cerqueira-Silva T, Pescarini JM, Cardim LL, Leyrat C, Whitaker H, Antunes de Brito CA, et al. Risk of death following chikungunya virus disease in the 100 Million Brazilian Cohort, 2015–18: a matched cohort study and self-controlled case series. The Lancet Infectious Diseases. 2024;24(5):504–13. doi: 10.1016/S1473-3099(23)00739-9.

6. Brito CAAd. Alert: Severe cases and deaths associated with Chikungunya in Brazil. Revista da Sociedade Brasileira de Medicina Tropical. 2017;50.

7. Rama K, de Roo AM, Louwsma T, Hofstra HS, Gurgel do Amaral GS, Vondeling GT, et al. Clinical outcomes of chikungunya: A systematic literature review and meta-analysis. PLOS Neglected Tropical Diseases. 2024;18(6):e0012254. doi: 10.1371/journal.pntd.0012254.

8. Brito C, Falcão MB, de Albuquerque M, Cerqueira-Silva T, Teixeira MG, Franca RFO. Chikungunya: From Hypothesis to Evidence of Increased Severe Disease and Fatalities. Viruses. 2025;17(1). Epub 2025/01/25. doi: 10.3390/v17010062. PubMed PMID: 39861851; PubMed Central PMCID: PMCPMC11768798.

9. McCarthy MK, Davenport BJJ, Morrison TE. Chronic Chikungunya Virus Disease. In: Heise M, editor. Chikungunya Virus. Cham: Springer International Publishing; 2022. p. 55–80.

10. Richardson JS, Anderson DM, Mendy J, Tindale LC, Muhammad S, Loreth T, et al. Chikungunya virus virus-like particle vaccine safety and immunogenicity in adolescents and adults in the USA: a phase 3, randomised, double-blind, placebo-controlled trial. The Lancet. 2025;405(10487):1343–52. doi: 10.1016/S0140-6736(25)00345-9.

11. Tindale LC, Richardson JS, Anderson DM, Mendy J, Muhammad S, Loreth T, et al. Chikungunya virus virus-like particle vaccine safety and immunogenicity in adults older than 65 years: a phase 3, randomised, double-blind, placebo-controlled trial. The Lancet. 2025;405(10487):1353–61. doi: 10.1016/S0140-6736(25)00372-1.

12. Schneider M, Narciso-Abraham M, Hadl S, McMahon R, Toepfer S, Fuchs U, et al. Safety and immunogenicity of a single-shot live-attenuated chikungunya vaccine: a double-blind, multicentre, randomised, placebo-controlled, phase 3 trial. The Lancet. 2023;401(10394):2138–47. doi: 10.1016/S0140-6736(23)00641-4.

13. The Lancet Infectious D. More data needed on the FDA’s decision to suspend Ixchiq. The Lancet Infectious Diseases. 2025;25(10):1055. doi: 10.1016/S1473-3099(25)00552-3.

14. Zhao J, Andreev I, Silva HM. Resident tissue macrophages: Key coordinators of tissue homeostasis beyond immunity. Science Immunology. 2024;9(94):eadd1967. doi: doi:10.1126/sciimmunol.add1967.

15. Gardner J, Anraku I, Le TT, Larcher T, Major L, Roques P, et al. Chikungunya virus arthritis in adult wild-type mice. J Virol. 2010;84(16):8021–32. Epub 2010/06/04. doi: 10.1128/jvi.02603-09. PubMed PMID: 20519386; PubMed Central PMCID: PMCPMC2916516.

16. Morrison Thomas E, Whitmore Alan C, Shabman Reed S, Lidbury Brett A, Mahalingam S, Heise Mark T. Characterization of Ross River Virus Tropism and Virus-Induced Inflammation in a Mouse Model of Viral Arthritis and Myositis. Journal of Virology. 2006;80(2):737–49. doi: 10.1128/jvi.80.2.737-749.2006.

17. Hawman DW, Stoermer KA, Montgomery SA, Pal P, Oko L, Diamond MS, et al. Chronic joint disease caused by persistent Chikungunya virus infection is controlled by the adaptive immune response. J Virol. 2013;87(24):13878–88. Epub 2013/10/18. doi: 10.1128/jvi.02666-13. PubMed PMID: 24131709; PubMed Central PMCID: PMCPMC3838294.

18. Zarrella KM, Sheridan RM, Ware BC, Davenport BJ, da Silva MOL, Vyshenska D, et al. Chikungunya virus persists in joint-associated macrophages and promotes chronic disease. Res Sq. 2025. Epub 2025/07/18. doi: 10.21203/rs.3.rs-6917990/v1. PubMed PMID: 40678223; PubMed Central PMCID: PMCPMC12270208 exists. La Jolla Institute for Immunology has filed for patent protection for various aspects of T cell epitope and vaccine design work.

19. Labadie K, Larcher T, Joubert C, Mannioui A, Delache B, Brochard P, et al. Chikungunya disease in nonhuman primates involves long-term viral persistence in macrophages. J Clin Invest. 2010;120(3):894–906. Epub 2010/02/25. doi: 10.1172/jci40104. PubMed PMID: 20179353; PubMed Central PMCID: PMCPMC2827953.

20. Hoarau J-J, Jaffar Bandjee M-C, Krejbich Trotot P, Das T, Li-Pat-Yuen G, Dassa B, et al. Persistent Chronic Inflammation and Infection by Chikungunya Arthritogenic Alphavirus in Spite of a Robust Host Immune Response. The Journal of Immunology. 2010;184(10):5914–27. doi: 10.4049/jimmunol.0900255.

21. Sharp TM, Keating MK, Shieh W-J, Bhatnagar J, Bollweg BC, Levine R, et al. Clinical Characteristics, Histopathology, and Tissue Immunolocalization of Chikungunya Virus Antigen in Fatal Cases. Clinical Infectious Diseases. 2021;73(2):e345–e54. doi: 10.1093/cid/ciaa837.

22. Holmes AC, Lucas CJ, Brisse ME, Ware BC, Hickman HD, Morrison TE, et al. Ly6C^+^ monocytes in the skin promote systemic alphavirus dissemination. Cell Reports. 2024;43(3). doi: 10.1016/j.celrep.2024.113876.

23. Lum F-M, Ng LFP. Cellular and molecular mechanisms of chikungunya pathogenesis. Antiviral Research. 2015;120:165–74. doi: 10.1016/j.antiviral.2015.06.009.

24. Valdés-López JF, Fernandez GJ, Urcuqui-Inchima S. Synergistic Effects of Toll-Like Receptor 1/2 and Toll-Like Receptor 3 Signaling Triggering Interleukin 27 Gene Expression in Chikungunya Virus-Infected Macrophages. Frontiers in Cell and Developmental Biology. 2022;10. doi: 10.3389/fcell.2022.812110.

25. Valdés-López JF, Fernandez GJ, Urcuqui-Inchima S. Interleukin 27 as an inducer of antiviral response against chikungunya virus infection in human macrophages. Cellular Immunology. 2021;367:104411. doi: 10.1016/j.cellimm.2021.104411.

26. Felipe VLJ, Paula A V, Silvio U-I. Chikungunya virus infection induces differential inflammatory and antiviral responses in human monocytes and monocyte-derived macrophages. Acta Tropica. 2020;211:105619. doi: 10.1016/j.actatropica.2020.105619.

27. Sourisseau M, Schilte C, Casartelli N, Trouillet C, Guivel-Benhassine F, Rudnicka D, et al. Characterization of Reemerging Chikungunya Virus. PLOS Pathogens. 2007;3(6):e89. doi: 10.1371/journal.ppat.0030089.

28. Trus E, Basta S, Gee K. Who’s in charge here? Macrophage colony stimulating factor and granulocyte macrophage colony stimulating factor: Competing factors in macrophage polarization. Cytokine. 2020;127:154939. doi: 10.1016/j.cyto.2019.154939.

29. Becher B, Tugues S, Greter M. GM-CSF: From Growth Factor to Central Mediator of Tissue Inflammation. Immunity. 2016;45(5):963–73. doi: 10.1016/j.immuni.2016.10.026.

30. de Souza WM, Fumagalli MJ, de Lima STS, Parise PL, Carvalho DCM, Hernandez C, et al. Pathophysiology of chikungunya virus infection associated with fatal outcomes. Cell Host & Microbe. 2024;32(4):606–22.e8. doi: 10.1016/j.chom.2024.02.011.

31. Chow A, Her Z, Ong EKS, Chen J-m, Dimatatac F, Kwek DJC, et al. Persistent Arthralgia Induced by Chikungunya Virus Infection is Associated with Interleukin-6 and Granulocyte Macrophage Colony-Stimulating Factor. The Journal of Infectious Diseases. 2011;203(2):149–57. doi: 10.1093/infdis/jiq042.

32. Mun SH, Park PSU, Park-Min K-H. The M-CSF receptor in osteoclasts and beyond. Experimental & Molecular Medicine. 2020;52(8):1239–54. doi: 10.1038/s12276-020-0484-z.

33. Wauquier N, Becquart P, Nkoghe D, Padilla C, Ndjoyi-Mbiguino A, Leroy EM. The acute phase of Chikungunya virus infection in humans is associated with strong innate immunity and T CD8 cell activation. J Infect Dis. 2011;204(1):115–23. Epub 2011/06/02. doi: 10.1093/infdis/jiq006. PubMed PMID: 21628665; PubMed Central PMCID: PMCPMC3307152.

34. Ushach I, Zlotnik A. Biological role of granulocyte macrophage colony-stimulating factor (GM-CSF) and macrophage colony-stimulating factor (M-CSF) on cells of the myeloid lineage. Journal of Leukocyte Biology. 2016;100(3):481–9. doi: 10.1189/jlb.3RU0316-144R.

35. Lum F-M, Chan Y-H, Teo T-H, Becht E, Amrun SN, Teng KWW, et al. Crosstalk between CD64^+^MHCII^+^ macrophages and CD4^+^ T cells drives joint pathology during chikungunya. EMBO Molecular Medicine. 2024;16(3):641–63-63. doi: 10.1038/s44321-024-00028-y.

36. De Luca G, Cavalli G, Campochiaro C, Della-Torre E, Angelillo P, Tomelleri A, et al. GM-CSF blockade with mavrilimumab in severe COVID-19 pneumonia and systemic hyperinflammation: a single-centre, prospective cohort study. The Lancet Rheumatology. 2020;2(8):e465–e73. doi: 10.1016/S2665-9913(20)30170-3.

37. Pupim L, Wang TS, Hudock K, Denson J, Fourie N, Hercilla Vasquez L, et al. LB0001 MAVRILIMUMAB IMPROVES OUTCOMES IN PHASE 2 TRIAL IN NON-MECHANICALLY-VENTILATED PATIENTS WITH SEVERE COVID-19 PNEUMONIA AND SYSTEMIC HYPERINFLAMMATION. Annals of the Rheumatic Diseases. 2021;80:198–9. doi: 10.1136/annrheumdis-2021-eular.5012.

38. Wang H, Tumes DJ, Hercus TR, Yip KH, Aloe C, Vlahos R, et al. Blocking the human common beta subunit of the GM-CSF, IL-5 and IL-3 receptors markedly reduces hyperinflammation in ARDS models. Cell Death & Disease. 2022;13(2):137. doi: 10.1038/s41419-022-04589-z.

39. Bouzeineddine NZ, Philippi A, Gee K, Basta S. Granulocyte macrophage colony stimulating factor in virus-host interactions and its implication for immunotherapy. Cytokine & Growth Factor Reviews. 2025;81:54–63. doi: 10.1016/j.cytogfr.2024.12.002.

40. García-Domínguez M. Targeting GM-CSF in Rheumatoid Arthritis: Advances in Cytokine-Directed Immunotherapy and Clinical Implications. Life [Internet]. 2025; 15(11):[1737 p.].

41. Wang Q, Wang J, Xu K, Luo Z. Targeting the CSF1/CSF1R signaling pathway: an innovative strategy for ultrasound combined with macrophage exhaustion in pancreatic cancer therapy. Front Immunol. 2024;15:1481247. Epub 2024/10/17. doi: 10.3389/fimmu.2024.1481247. PubMed PMID: 39416792; PubMed Central PMCID: PMCPMC11479911.

42. Miner JJ, Aw Yeang HX, Fox JM, Taffner S, Malkova ON, Oh ST, et al. Brief Report: Chikungunya Viral Arthritis in the United States: A Mimic of Seronegative Rheumatoid Arthritis. Arthritis & Rheumatology. 2015;67(5):1214–20. doi: 10.1002/art.39027.

43. Stoermer KA, Burrack A, Oko L, Montgomery SA, Borst LB, Gill RG, et al. Genetic ablation of arginase 1 in macrophages and neutrophils enhances clearance of an arthritogenic alphavirus. J Immunol. 2012;189(8):4047–59. Epub 2012/09/14. doi: 10.4049/jimmunol.1201240. PubMed PMID: 22972923; PubMed Central PMCID: PMCPMC3466331.

44. Burrack KS, Tan JJ, McCarthy MK, Her Z, Berger JN, Ng LF, et al. Myeloid Cell Arg1 Inhibits Control of Arthritogenic Alphavirus Infection by Suppressing Antiviral T Cells. PLoS Pathog. 2015;11(10):e1005191. Epub 2015/10/06. doi: 10.1371/journal.ppat.1005191. PubMed PMID: 26436766; PubMed Central PMCID: PMCPMC4593600.

45. Hernández-Sarmiento LJ, Tamayo-Molina YS, Valdés-López JF, Urcuqui-Inchima S. Mayaro virus infection elicits a robust pro-inflammatory and antiviral response in human macrophages. Acta Tropica. 2024;252:107146. doi: 10.1016/j.actatropica.2024.107146.

46. Kril V, Aïqui-Reboul-Paviet O, Briant L, Amara A. New Insights into Chikungunya Virus Infection and Pathogenesis. Annual Review of Virology. 2021;8(Volume 8, 2021):327–47. doi: 10.1146/annurev-virology-091919-102021.

47. Noval MG, Spector SN, Bartnicki E, Izzo F, Narula N, Yeung ST, et al. MAVS signaling is required for preventing persistent chikungunya heart infection and chronic vascular tissue inflammation. Nature Communications. 2023;14(1):4668. doi: 10.1038/s41467-023-40047-w.

48. Berry GE, Tse LV. Virus Binding and Internalization Assay for Adeno-associated Virus. Bio Protoc. 2017;7(2). Epub 2017/12/12. doi: 10.21769/BioProtoc.2110. PubMed PMID: 29226179; PubMed Central PMCID: PMCPMC5718205.

49. Zimmerman O, Holmes AC, Kafai NM, Adams LJ, Diamond MS. Entry receptors - the gateway to alphavirus infection. J Clin Invest. 2023;133(2). Epub 2023/01/18. doi: 10.1172/jci165307. PubMed PMID: 36647825; PubMed Central PMCID: PMCPMC9843064 Biosciences, Merck, Novavax, Moderna, and Immunome. The Diamond laboratory has received unrelated funding support in sponsored research agreements from Vir Biotechnology, Moderna, Immunome, and Emergent BioSolutions.

50. Mercer J, Greber UF. Virus interactions with endocytic pathways in macrophages and dendritic cells. Trends in Microbiology. 2013;21(8):380–8. doi: 10.1016/j.Vm.2013.06.001.

51. Bernard E, Solignat M, Gay B, Chazal N, Higgs S, Devaux C, et al. Endocytosis of Chikungunya Virus into Mammalian Cells: Role of Clathrin and Early Endosomal Compartments. PLOS ONE. 2010;5(7):e11479. doi: 10.1371/journal.pone.0011479.

52. Mylvaganam S, Freeman SA, Grinstein S. The cytoskeleton in phagocytosis and macropinocytosis. Current Biology. 2021;31(10):R619–R32. doi: 10.1016/j.cub.2021.01.036.

53. Schulz D, Severin Y, Zanotelli VRT, Bodenmiller B. In-Depth Characterization of Monocyte-Derived Macrophages using a Mass Cytometry-Based Phagocytosis Assay. Scientific Reports. 2019;9(1):1925. doi: 10.1038/s41598-018-38127-9.

54. Lee CHR, Mohamed Hussain K, Chu JJH. Macropinocytosis dependent entry of Chikungunya virus into human muscle cells. PLOS Neglected Tropical Diseases. 2019;13(8):e0007610. doi: 10.1371/journal.pntd.0007610.

55. Hoornweg Tabitha E, van Duijl-Richter Mareike KS, Ayala Nuñez Nilda V, Albulescu Irina C, van Hemert Martijn J, Smit Jolanda M. Dynamics of Chikungunya Virus Cell Entry Unraveled by Single-Virus Tracking in Living Cells. Journal of Virology. 2016;90(9):4745–56. doi: 10.1128/jvi.03184-15.

56. Gold ES, Underhill DM, MorrisseCe NS, Guo J, McNiven MA, Aderem A. Dynamin 2 Is Required for Phagocytosis in Macrophages. Journal of Experimental Medicine. 1999;190(12):1849–56. doi: 10.1084/jem.190.12.1849.

57. Smith SM, Smith CJ. Capturing the mechanics of clathrin-mediated endocytosis. Current Opinion in Structural Biology. 2022;75:102427. doi: 10.1016/j.sbi.2022.102427.

58. De Caluwé L, Ariën KK, Bartholomeeusen K. Host Factors and Pathways Involved in the Entry of Mosquito-Borne Alphaviruses. Trends in Microbiology. 2021;29(7):634–47. doi: 10.1016/j.Vm.2020.10.011.

59. Yao Z, Ramachandran S, Huang S, Kim E, Jami-Alahmadi Y, Kaushal P, et al. Interaction of chikungunya virus glycoproteins with macrophage factors controls virion production. The EMBO Journal. 2024;43(20):4625–55-55. doi: 10.1038/s44318-024-00193-3.

60. Jemielity S, Wang JJ, Chan YK, Ahmed AA, Li W, Monahan S, et al. TIM-family Proteins Promote Infection of Multiple Enveloped Viruses through Virion-associated Phosphatidylserine. PLOS Pathogens. 2013;9(3):e1003232. doi: 10.1371/journal.ppat.1003232.

61. Li FS, Carpentier KS, Hawman DW, Lucas CJ, Ander SE, Feldmann H, et al. Species-specific MARCO-alphavirus interactions dictate chikungunya virus viremia. Cell Rep. 2023;42(5):112418. Epub 2023/04/21. doi: 10.1016/j.celrep.2023.112418. PubMed PMID: 37083332; PubMed Central PMCID: PMCPMC10290254.

62. Zhang R, Kim AS, Fox JM, Nair S, Basore K, Klimstra WB, et al. Mxra8 is a receptor for multiple arthritogenic alphaviruses. Nature. 2018;557(7706):570–4. doi: 10.1038/s41586-018-0121-3.

63. Samaniego R, Palacios BS, Domiguez-Soto Á, Vidal C, Salas A, Matsuyama T, et al. Macrophage uptake and accumulation of folates are polarization-dependent in vitro and in vivo and are regulated by activin A. Journal of Leukocyte Biology. 2014;95(5):797–808. doi: 10.1189/jlb.0613345.

64. Kruger M, Van De Winkel JGJ, De Wit TPM, Coorevits L, Ceuppens JL. Granulocyte–macropha ge colony-stimulating factor down-regulates CD14 expression on monocytes. Immunology. 1996;89(1):89–95. doi: 10.1046/j.1365-2567.1996.d01-707.x.

65. Baumann CL, Aspalter IM, Sharif O, Pichlmair A, Blüml S, Grebien F, et al. CD14 is a coreceptor of Toll-like receptors 7 and 9. J Exp Med. 2010;207(12):2689–701. Epub 2010/11/17. doi: 10.1084/jem.20101111. PubMed PMID: 21078886; PubMed Central PMCID: PMCPMC2989773.

66. Reis e Sousa C, Yamasaki S, Brown GD. Myeloid C-type lectin receptors in innate immune recognition. Immunity. 2024;57(4):700–17. doi: 10.1016/j.immuni.2024.03.005.

67. van der Zande HJP, Nitsche D, Schlautmann L, Guigas B, Burgdorf S. The Mannose Receptor: From Endocytic Receptor and Biomarker to Regulator of (Meta)Inflammation. Frontiers in Immunology. 2021;Volume 12 - 2021. doi: 10.3389/fimmu.2021.765034.

68. Conforti A, Lavalle G, Varini F, LuccheÇ L, Cataldi G, Faticoni A, et al. Pro-Inflammatory Cytokines as Early Predictors of Chronic Rheumatologic Disease Following Chikungunya Virus Infection. J Clin Med. 2025;14(19). Epub 2025/10/16. doi: 10.3390/jcm14196720. PubMed PMID: 41095803; PubMed Central PMCID: PMCPMC12525192.

69. Kayesh MEH, Kohara M, Tsukiyama-Kohara K. Toll-like Receptor (TLR) Response in Chikungunya Virus Infection: Mechanism of Activation, Immune Evasion, and Use of TLR Agonists in Vaccine Development. Vaccines (Basel). 2025;13(8). Epub 2025/08/30. doi: 10.3390/vaccines13080856. PubMed PMID: 40872941; PubMed Central PMCID: PMCPMC12390013.

70. Webb LG, Veloz J, Pintado-Silva J, Zhu T, Rangel MV, Mutetwa T, et al. Chikungunya virus antagonizes cGAS-STING mediated type-I interferon responses by degrading cGAS. PLOS Pathogens. 2020;16(10):e1008999. doi: 10.1371/journal.ppat.1008999.

71. Fros Jelke J, Liu Wen J, Prow Natalie A, Geertsema C, Ligtenberg M, Vanlandingham Dana L, et al. Chikungunya Virus Nonstructural Protein 2 Inhibits Type I/II Interferon-Stimulated JAK-STAT Signaling. Journal of Virology. 2010;84(20):10877–87. doi: 10.1128/jvi.00949-10.

72. Pott F, Postmus D, Brown RJP, Wyler E, Neumann E, Landthaler M, et al. Single-cell analysis of arthritogenic alphavirus-infected human synovial fibroblasts links low abundance of viral RNA to induction of innate immunity and arthralgia-associated gene expression. Emerg Microbes Infect. 2021;10(1):2151–68. Epub 2021/11/02. doi: 10.1080/22221751.2021.2000891. PubMed PMID: 34723780; PubMed Central PMCID: PMCPMC8604527.

73. Werneke SW, Schilte C, Rohatgi A, Monte KJ, Michault A, Arenzana-Seisdedos F, et al. ISG15 is critical in the control of Chikungunya virus infection independent of UbE1L mediated conjugation. PLoS Pathog. 2011;7(10):e1002322. Epub 2011/10/27. doi: 10.1371/journal.ppat.1002322. PubMed PMID: 22028657; PubMed Central PMCID: PMCPMC3197620.

74. Rodriguez Marisela R, Monte K, Thackray Larissa B, Lenschow Deborah J. ISG15 Functions as an Interferon-Mediated Antiviral Effector Early in the Murine Norovirus Life Cycle. Journal of Virology. 2014;88(16):9277–86. doi: 10.1128/jvi.01422-14.

75. Villanueva Guzman MD, Yao Z, Li MMH, Noval MG. Hidden in Plain Sight: Alphavirus Persistence and Its Potential for Driving Chronic Pathogenesis. Viruses [Internet]. 2026; 18(1):[30 p.].

76. Choi Y, Bowman JW, Jung JU. Autophagy during viral infection — a double-edged sword. Nature Reviews Microbiology. 2018;16(6):341–54. doi: 10.1038/s41579-018-0003-6.

77. Zhang Y, Morgan MJ, Chen K, Choksi S, Liu Z-g. Induction of autophagy is essential for monocyte-macrophage differentiation. Blood. 2012;119(12):2895–905. doi: 10.1182/blood-2011-08-372383.

78. Atri C, Guerfali FZ, Laouini D. Role of Human Macrophage Polarization in Inflammation during Infectious Diseases. Int J Mol Sci. 2018;19(6). Epub 2018/06/21. doi: 10.3390/ijms19061801. PubMed PMID: 29921749; PubMed Central PMCID: PMCPMC6032107.

79. Wang Y, Zhao M, Liu S, Guo J, Lu Y, Cheng J, et al. Macrophage-derived extracellular vesicles: diverse mediators of pathology and therapeutics in multiple diseases. Cell Death & Disease. 2020;11(10):924. doi: 10.1038/s41419-020-03127-z.

80. Upham Jacqueline P, Pickett D, Irimura T, Anders EM, Reading Patrick C. Macrophage Receptors for Influenza A Virus: Role of the Macrophage Galactose-Type Lectin and Mannose Receptor in Viral Entry. Journal of Virology. 2010;84(8):3730–7. doi: 10.1128/jvi.02148-09.

81. Wu MF, Chen ST, Yang AH, Lin WW, Lin YL, Chen NJ, et al. CLEC5A is critical for dengue virus-induced inflammasome activation in human macrophages. Blood. 2013;121(1):95–106. Epub 2012/11/16. doi: 10.1182/blood-2012-05-430090. PubMed PMID: 23152543.

82. Lv J, Wang Z, Qu Y, Zhu H, Zhu Q, Tong W, et al. Distinct uptake, amplification, and release of SARS-CoV-2 by M1 and M2 alveolar macrophages. Cell Discovery. 2021;7(1):24. doi: 10.1038/s41421-021-00258-1.

83. Patel AR, Garg A, Rosberger HT, Kowdle S, Reis RA, Frere JJ, et al. Alveolar macrophages are early targets of mumps virus. Proceedings of the National Academy of Sciences. 2024;121(52):e2410954121. doi: 10.1073/pnas.2410954121.

84. dos Santos CF, Nunes PCG, Fiestas-Solorzano VE, Gandini M, dos Santos FB, Pinheiro RO, et al. Monocyte Dynamics in Chikungunya Fever: Sustained Activation and Vascular-Coagulation Pathway Involvement. Viruses. 2025;17(9):1224. PubMed PMID: 10.3390/v17091224.

85. Stone AEL, Green R, Wilkins C, Hemann EA, Gale M, Jr. RIG-I-like receptors direct inflammatory macrophage polarization against West Nile virus infection. Nat Commun. 2019;10(1):3649. Epub 2019/08/15. doi: 10.1038/s41467-019-11250-5. PubMed PMID: 31409781; PubMed Central PMCID: PMCPMC6692387.

86. Fleetwood AJ, Dinh H, Cook AD, Hertzog PJ, Hamilton JA. GM-CSF- and M-CSF-dependent macrophage phenotypes display differential dependence on Type I interferon signaling. Journal of Leukocyte Biology. 2009;86(2):411–21. doi: 10.1189/jlb.1108702.

87. Rodriguez RM, Suarez-Alvarez B, Lavín JL, Ascensión AM, Gonzalez M, Lozano JJ, et al. Signal Integration and Transcriptional Regulation of the Inflammatory Response Mediated by the GM-/M-CSF Signaling Axis in Human Monocytes. Cell Reports. 2019;29(4):860–72.e5. doi: 10.1016/j.celrep.2019.09.035.

88. Simón-Fuentes M, Herrero C, Acero-Riaguas L, Nieto C, Lasala F, Labiod N, et al. TLR7 Activation in M-CSF-Dependent Monocyte-Derived Human Macrophages Potentiates Inflammatory Responses and Prompts Neutrophil Recruitment. Journal of Innate Immunity. 2023;15(1):517–30. doi: 10.1159/000530249.

89. Her Z, Teng TS, Tan JJL, Teo TH, Kam YW, Lum FM, et al. Loss of TLR3 aggravates CHIKV replication and pathology due to an altered virus-specific neutralizing antibody response. EMBO Molecular Medicine. 2015;7(1):24–41. doi: 10.15252/emmm.201404459.

90. Lundberg AM, Drexler SK, Monaco C, Williams LM, Sacre SM, Feldmann M, et al. Key differences in TLR3/poly I:C signaling and cytokine induction by human primary cells: a phenomenon absent from murine cell systems. Blood. 2007;110(9):3245–52. 10.1182/blood-2007-02-072934.

91. Sadeghi K, Wisgrill L, Wessely I, Diesner SC, Schüller S, Dürr C, et al. GM-CSF Down-Regulates TLR Expression via the Transcription Factor PU.1 in Human Monocytes. PLoS One. 2016;11(10):e0162667. Epub 2016/10/04. doi: 10.1371/journal.pone.0162667. PubMed PMID: 27695085; PubMed Central PMCID: PMCPMC5047522.

92. Mansell A, Smith R, Doyle SL, Gray P, Fenner JE, Crack PJ, et al. Suppressor of cytokine signaling 1 negatively regulates Toll-like receptor signaling by mediating Mal degradation. Nature Immunology. 2006;7(2):148–55. doi: 10.1038/ni1299.

93. Whyte CS, Bishop ET, Rückerl D, Gaspar-Pereira S, Barker RN, Allen JE, et al. Suppressor of cytokine signaling (SOCS)1 is a key determinant of differential macrophage activation and function. Journal of Leukocyte Biology. 2011;90(5):845–54. doi: 10.1189/jlb.1110644.

94. Liu S, Wang M, Xu L, Deng D, Lu L, Tian J, et al. New insight into the role of SOCS family in immune regulation and autoimmune pathogenesis. Journal of Advanced Research. 2025. doi: 10.1016/j.jare.2025.05.020.

95. Bunda S, Kommaraju K, Heir P, Ohh M. SOCS-1 Mediates Ubiquitylation and Degradation of GM-CSF Receptor. PLOS ONE. 2013;8(9):e76370. doi: 10.1371/journal.pone.0076370.

96. Sobah ML, Liongue C, Ward AC. SOCS Proteins in Immunity, Inflammatory Diseases, and Immune-Related Cancer. Frontiers in Medicine. 2021;Volume 8 - 2021. doi: 10.3389/fmed.2021.727987.

97. Akhtar Lisa N, Benveniste ECy N. Viral Exploitation of Host SOCS Protein Functions. Journal of Virology. 2011;85(5):1912–21. doi: 10.1128/jvi.01857-10.

98. Low ZY, Wen Yip AJ, Chow VTK, Lal SK. The Suppressor of Cytokine Signalling family of proteins and their potential impact on COVID-19 disease progression. Reviews in Medical Virology. 2022;32(3):e2300. doi: 10.1002/rmv.2300.

99. Pintado Silva J, Fenutria R, Bernal-Rubio D, Sanchez-Martin I, Hunziker A, Chebishev E, et al. The dengue virus 4 component of NIAID’s tetravalent TV003 vaccine drives its innate immune signature. Experimental Biology and Medicine. 2022;247(24):2201–12. doi: 10.1177/15353702231151241.

