## Supplementary figures and images for "GM-CSF and M-CSF Driven Differentiation Differentially Regulates Chikungunya Virus Infection and Antiviral Responses in Human Monocyte-Derived Macrophages"

### S1 Fig

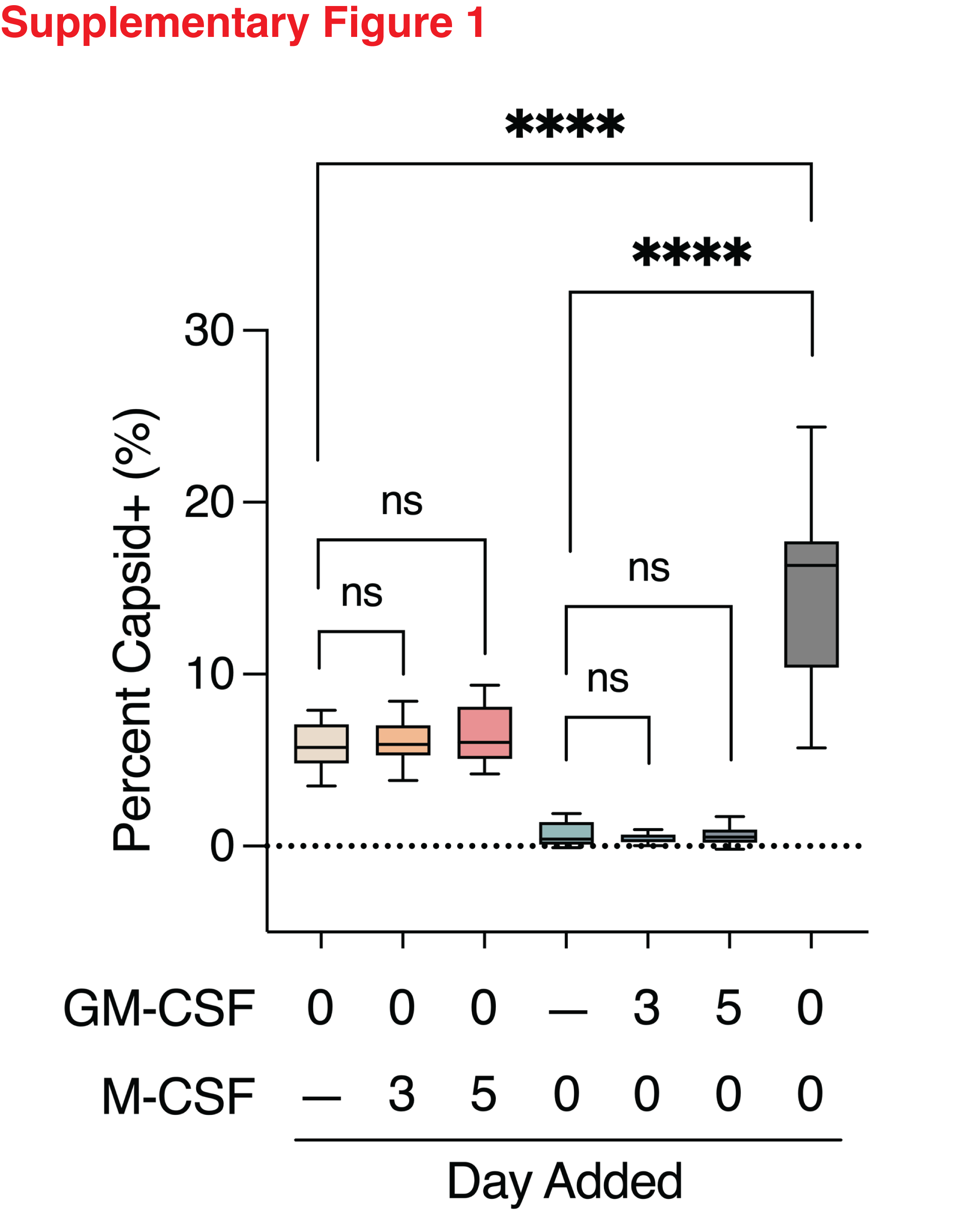

### S2 Fig

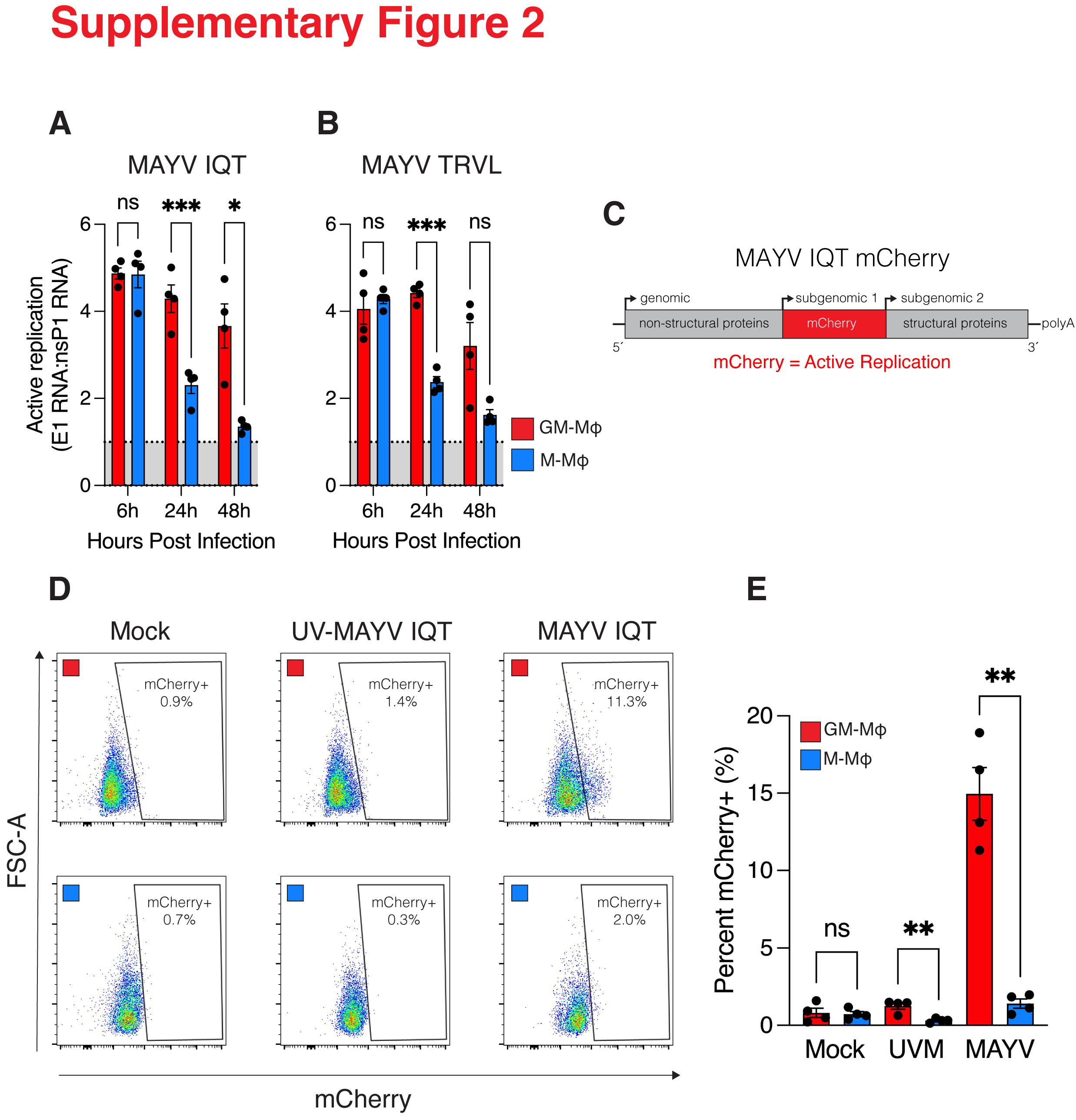

### S3 Fig

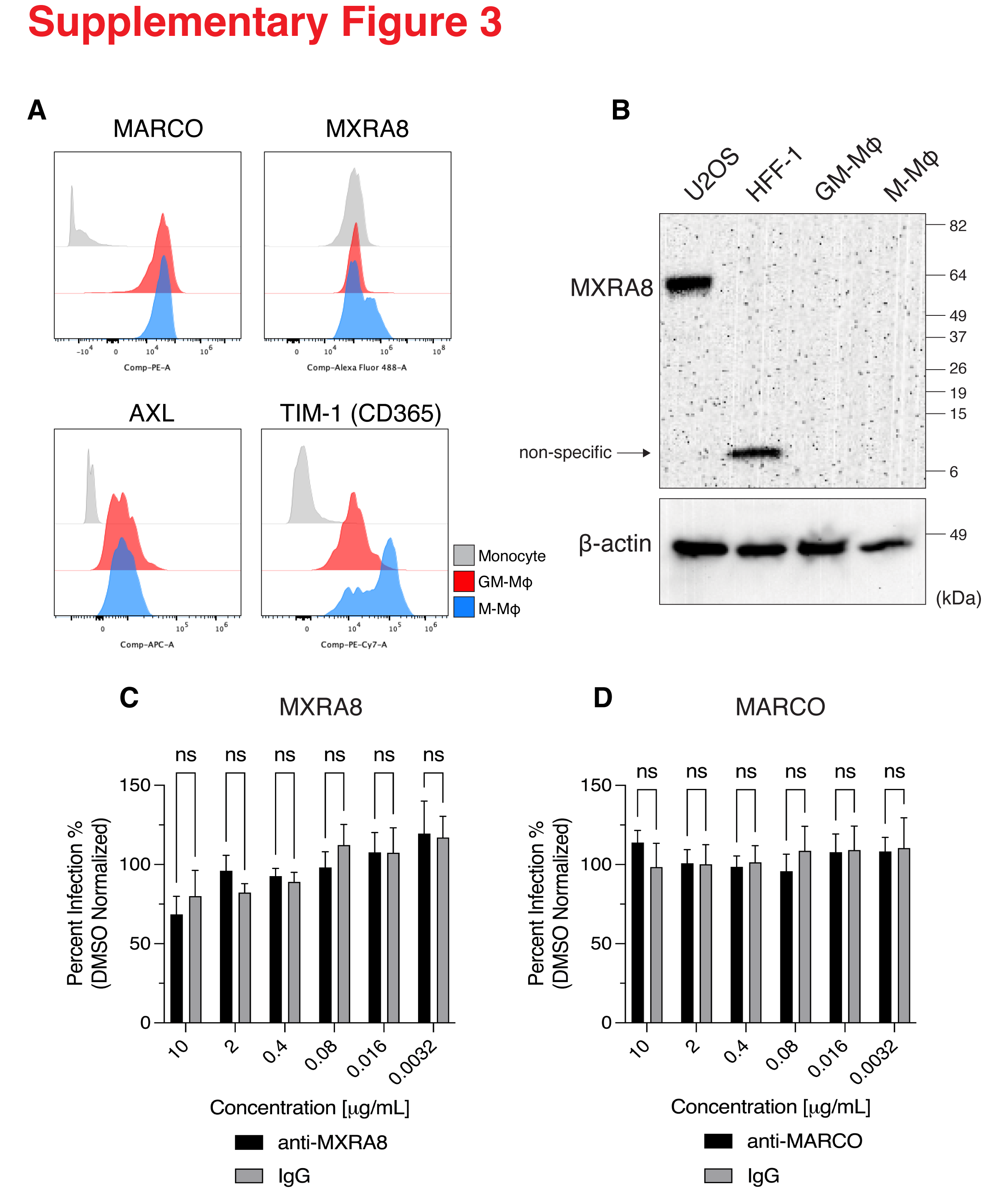

### S4 Fig

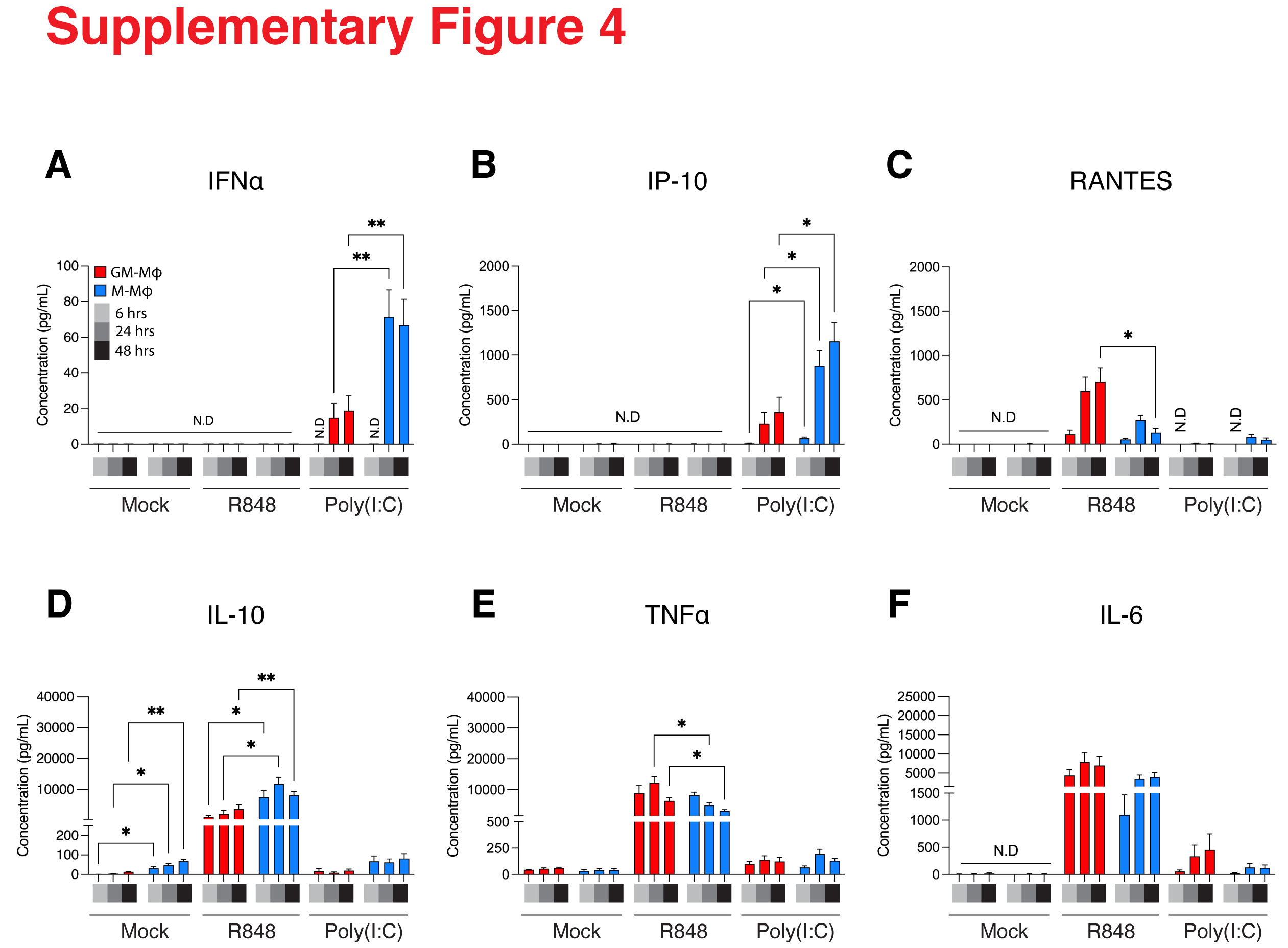
